# Structural basis of TRPV1 modulation by endogenous bioactive lipids

**DOI:** 10.1101/2023.05.11.540281

**Authors:** William R. Arnold, Adamo Mancino, Frank R. Moss, Adam Frost, David Julius, Yifan Cheng

**Author notes:** Correspondence: Yifan Cheng,; David Julius.

## Abstract

TRP ion channels are modulated by phosphoinositide lipids, but the underlying structural mechanisms remain unclear. The capsaicin- and heat-activated receptor, TRPV1, has served as a model for deciphering lipid modulation, which is relevant to understanding how pro-algesic agents enhance channel activity in the setting of inflammatory pain. Identification of a pocket within the TRPV1 transmembrane core has provided initial clues as to how phosphoinositide lipids bind to and regulate the channel. Here we show that this regulatory pocket can accommodate diverse lipid species, including the inflammatory lipid lysophosphatidic acid (LPA), whose actions are determined by their specific modes of binding. Furthermore, we show that an ‘empty pocket’ channel lacking an endogenous phosphoinositide lipid assumes an agonist-like state, even at low temperature, substantiating the concept that phosphoinositide lipids serve as negative TRPV1 modulators whose ejection from the binding pocket is a critical step towards activation by thermal or chemical stimuli.

## Introduction

Foundational studies of TRP channels in the *Drosophila* phototransduction pathway implicated phosphoinositide lipids as key modulatory agents^1,2^. Specifically, both genetic and electrophysiological studies have suggested that hydrolysis of PIP_2_ is a critical step connecting activation of PLC-coupled rhodopsin to channel gating. This theme has since been extended to many members of the vertebrate TRP channel family, where both positive and negative modulatory effects of phosphoinositide lipids have been proposed^3^. Although TRPV1 is not strictly a ‘receptor-operated’ channel, its sensitivity to heat and chemical agonists is enhanced by pro-algesic agents, such as bradykinin and nerve growth factor, that activate PLC-coupled metabotropic receptors, consistent with the idea that hydrolysis of phosphoinositides disinhibits the channel to promote gating^4^. Our previous findings support this model by showing that that the vanilloid binding pocket (VBP) is a critical regulatory site that harbors an endogenous phosphoinositide lipid when TRPV1 is in its inactive closed state, and that vanilloid agonists displace this lipid in the course of activating the channel^5,6^.

While physiological studies have shown that multiple phosphoinositide lipid species can inhibit TRPV1^7^, structural analyses have not provided definitive identification of the entity found in the VBP of the inactive channel. Furthermore, the idea that phosphoinositide lipids inhibit TRPV1 is inconsistent with the observation that soluble PIP_2_ analogues enhance, rather than inhibit, TRPV1 activation^8,9^. Also at issue is whether endogenous pro-algesic lipids bind to the same site and mediate their effects by displacing the resident PI lipid from the VBP^10^ as has been observed with exogenous lipophilic ligands such as capsaicin and other vanilloids.

Here, we address these questions using cryogenic electron microscopy (cryo-EM), which allows us to visualize the occupancy of the VBP and how this relates to the functional state of the channel. We captured several states of TRPV1 ranging from channels in which the VBP is empty to those bearing different bioactive lipids, including distinct phosphoinositide species, brominated phosphoinositide analogues, and a pro-algesic lipid, lysophosphatidic acid (LPA). Together, our data show how the VBP accommodates a range of bioactive lipids and how occupancy of the site corresponds to the structural status of the pore.

## Results and Discussion

### Ejecting the resident PI lipid favors the open state

To determine the effect of the resident PI lipid on channel gating, we devised a protocol to eject this lipid and render the binding pocket empty (Figure 1a). First, we used capsaicin to displace the resident lipid from detergent-solubilized channels. We then washed out the competing vanilloid followed by reconstitution of the channel into lipid nanodiscs of defined composition and devoid of phosphoinositide lipids. These samples were applied to grids held at 4° or 25°C (temperatures at which TRPV1 is normally closed) and then taken forward for cryo-EM analysis. The final maps were refined using C4 symmetry and resolved to 2.9 Å and 3.7 Å for the 4° and 25°C samples, respectively.

**Figure 1.**
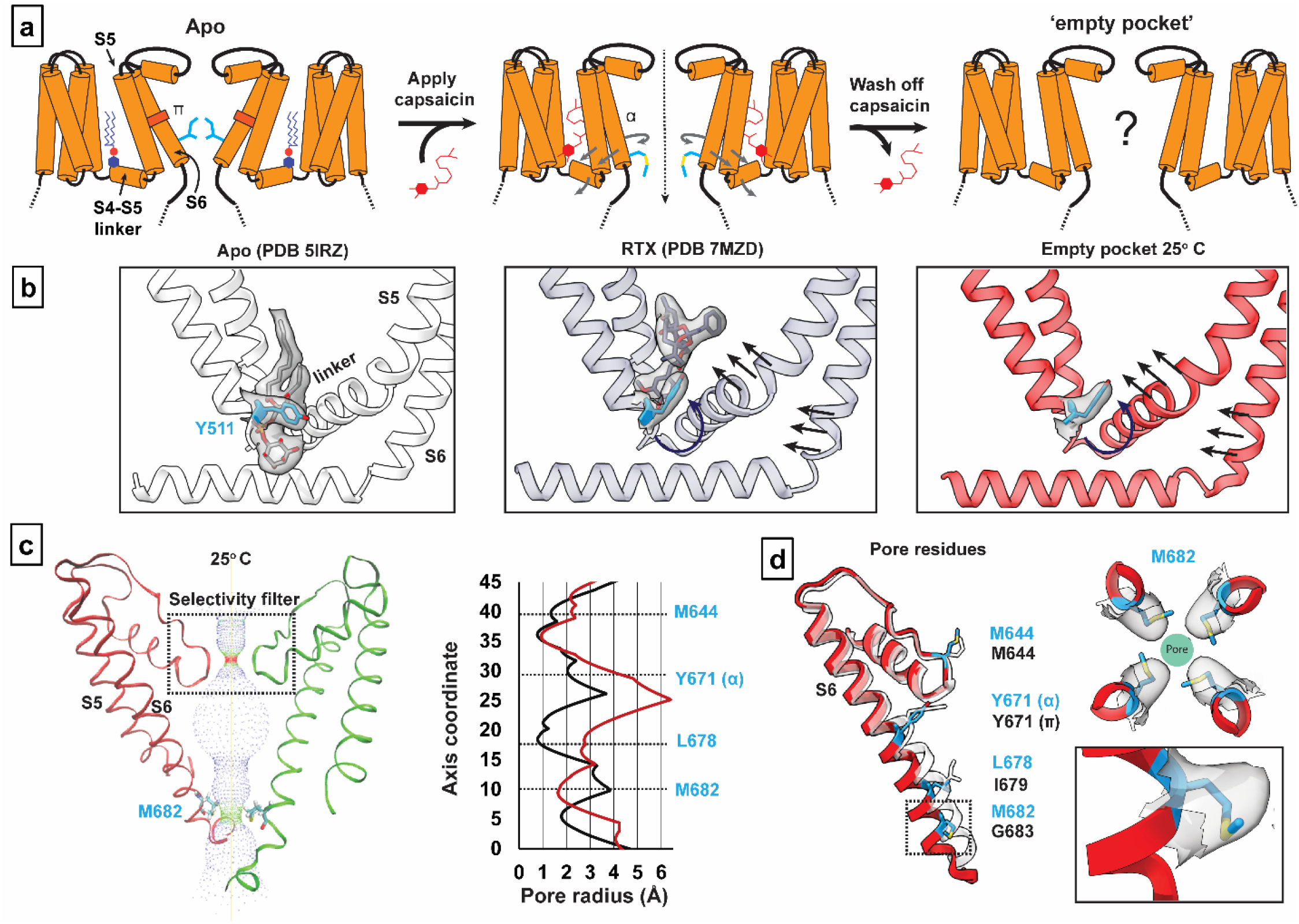
Empty-pocket TRPV1. **(a)** Schematic of the capsaicin washout procedure for obtaining empty-pocket TRPV1. **(b)** VBP in the apo state (PDB 5IRZ), bound with RTX (PDB 7MZD), or in the empty-pocket state at 25°C (PDB 8U3L; this study). **(c)** (left) Pore profile of empty-pocket TRPV1 at 25°C. (right) Pore radius of empty-pocket TRPV1 at 25°C (red) and apo TRPV1 (black) were determined using the HOLE program. **(d)** Key residues lining the channel pore. Red ribbon with blue labels depicts empty-pocket TRPV1 at 25°C; transparent ribbon represents apo TRPV1.

In both cryo-EM structures the vanilloid pocket is, indeed, unoccupied by either a resident PI lipid or capsaicin (Figure 1, Supplementary Figure 1). Previous functional and structural studies have identified tyrosine 511 (Y511) as a residue whose reorientation towards the vanilloid pocket correlates with ejection of the resident PI lipid upon ligand binding^5,6,11^. Interestingly, we observed that Y511 is flipped toward the binding pocket in these two structures, akin to what is seen in the agonist-bound state, demonstrating that reorientation of this fiducial side chain primarily reflects ejection of the resident lipid (Figure 1b). The empty space in the vanilloid pocket is now partially occupied by an acyl tail from an annular lipid in the outer leaflet (Supplementary Figure 1d) which does not reach the bottom of the pocket to restrict Y511 reorientation.

We next asked how these actions affect the ion permeation pathway. Overall, the pathway resembles that of vanilloid-activated channels in which the S4-S5 linker and S6 helices move away from the central axis (Figure 1c and 1d, Supplementary Figure 1a-1c). We also see transition of a π-helical turn in S6 to an α-helical configuration (π-α transition), resulting in rotation of the lower half of the S6 helix by one residue. This π-α transition rotates the gating residue L679 away from the central axis with a narrowest restriction now formed by rotation of M682 into the pore. This new restriction resembles that formed by M644 in the selectivity filter (Supplementary Figure 1e and 1f). Consistent with a previous study^6^, we see a presumptive Na^+^ ion coordinated to G643 in the selectivity filter, located just below M644, indicating that this configuration, in which methionine residues form the narrowest points in the ion permeation pathway, is conducting. Indeed, the B-factor surrounding M682 suggests that this residue is dynamic and thus sufficiently malleable to pass ions. Importantly, π-α transitions are seen when agonists, but not antagonists, occupy the VBP^5,6,11^, and these data now show that ejection of the resident lipid is also sufficient to support this transition. Interestingly, the main difference between 4° and 25° C structures was seen in the density of the M682 side chain, which was less well-resolved in the latter, implying a more dynamic nature at higher temperature. This contrasts with the π-α transition, which is seen at both temperatures and thus driven primarily by PI lipid ejection.

### PI and PIP_2_ favor the closed channel state

In all cryo-EM structures of TRPV1 reported to-date, the resident lipid shows clear features of a phosphoinositide, but the unresolved number of phosphate moieties on the inositol ring makes assignment to a specific species ambiguous. Moreover, the VBP is sufficiently flexible to accommodate diverse phosphoinositide species. Indeed, we have previously shown that phosphatidylinositol (PI) or phosphatidylinositol 4,5-bisphosphate (PIP_2_) inhibits TRPV1 when reconstituted into proteoliposomes^7^, suggesting that both species can bind to the VBP and stabilize a closed state. To resolve this question, we exploited our ability to generate TRPV1 protein with an empty vanilloid pocket, into which we introduced tetrabrominated analogues of PI or PIP_2_ (PI-Br_4_ and PIP_2_-Br_4_, respectively) as contrast-enhancing probes to distinguish these lipids from the other, unbrominated lipids in the nanodiscs (Figure 2a). Previous work has established that brominated lipids are faithful analogs of the unsaturated lipids from which they are synthesized^12^. Indeed, in these high-resolution structures (2.2-2.4 Å resolution) (Supplementary Figure 2) we were able to identify the bromine atoms within the oleoyl tails of the lipids by visually comparing the resulting reconstructions against those previously determined for samples containing endogenous lipids (Figure 2, Supplementary Figure 3). The well-resolved inositol headgroup of the bound lipids showed the expected differential location of phosphate moieties, most notably at the 4 and 5 positions for PIP_2_. Furthermore, conformations of both PI- and PIP_2_-reconstituted channels resemble the apo channel containing a native resident lipid. Together, these data demonstrate that the vanilloid site can accommodate PI or PIP_2_ where they act as negative regulatory factors to stabilize the closed state.

**Figure 2.**
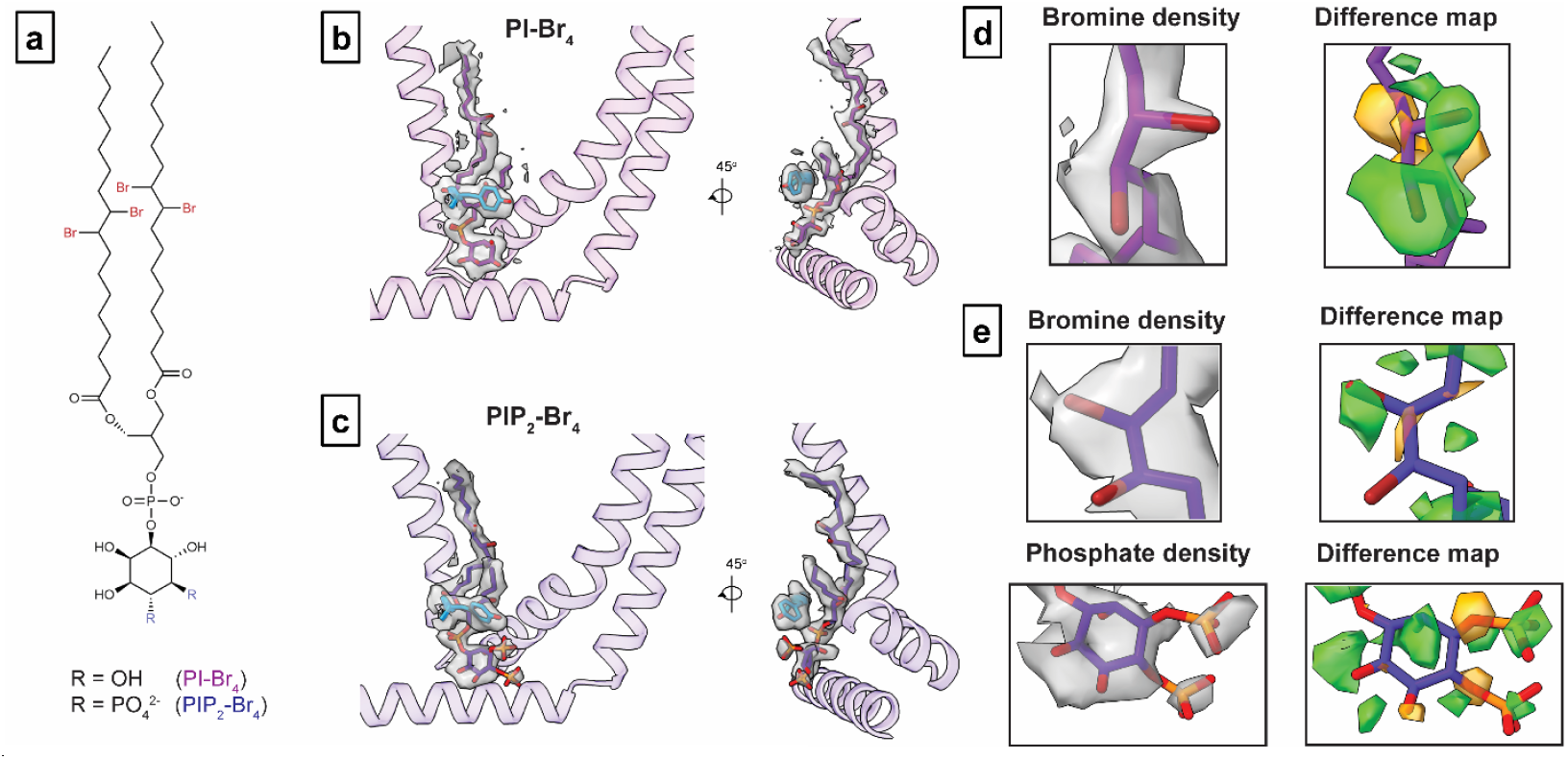
Brominated phospho-inositide binding to TRPV1. **(a)** Chemical structure of brominated phosphoinositides used in this study. **(b, c)** VBP with corresponding density for (b) PI-Br_4_ and (c) PIP_2_-Br_4_. **(d, e**) Density map of key functional groups for (d) PI-Br_4_ and (e) PIP_2_-Br_4_ with corresponding difference maps. Difference maps were determined by subtracting calculated map of models lacking bromine atoms and the phosphate moieties at 4 and 5 positions of the inositol headgroup from the experimental map (see methods). Both positive (green) and negative (orange) difference densities are shown.

### Soluble PIP_2_ is a partial TRPV1 potentiator

Physiological effects of phosphoinositides are often examined using soluble analogues of PIP_2_ with shortened acyl tails (such as diC8-PIP_2_) to facilitate cellular application during electrophysiological or imaging experiments. In the case of TRPV1, perfusion of diC8-PIP_2_ onto TRPV1-expressing cells has been shown to enhance channel activity, suggesting that phosphoinositide lipids serve as positive channel regulators^8,13^, conflicting with the idea that activation of PLC-coupled receptors potentiates TRPV1 activity through hydrolysis of PIP_2_ to the channel from phosphoinositide inhibition^4^. Considering this controversy, we were interested in determining whether soluble diC8-PIP_2_ and full-length PIP_2_ interact with TRPV1 in a similar manner. We therefore obtained cryo-EM data for samples in which diC8-PIP_2_ was provided to the ‘empty pocket’ TRPV1 preparation. Using focused classification, we were able to identify two key states, one corresponding to a closed state (3.0 Å) and the other having a dilated selectivity filter (3.6 Å) (Figure 3).

**Figure 3.**
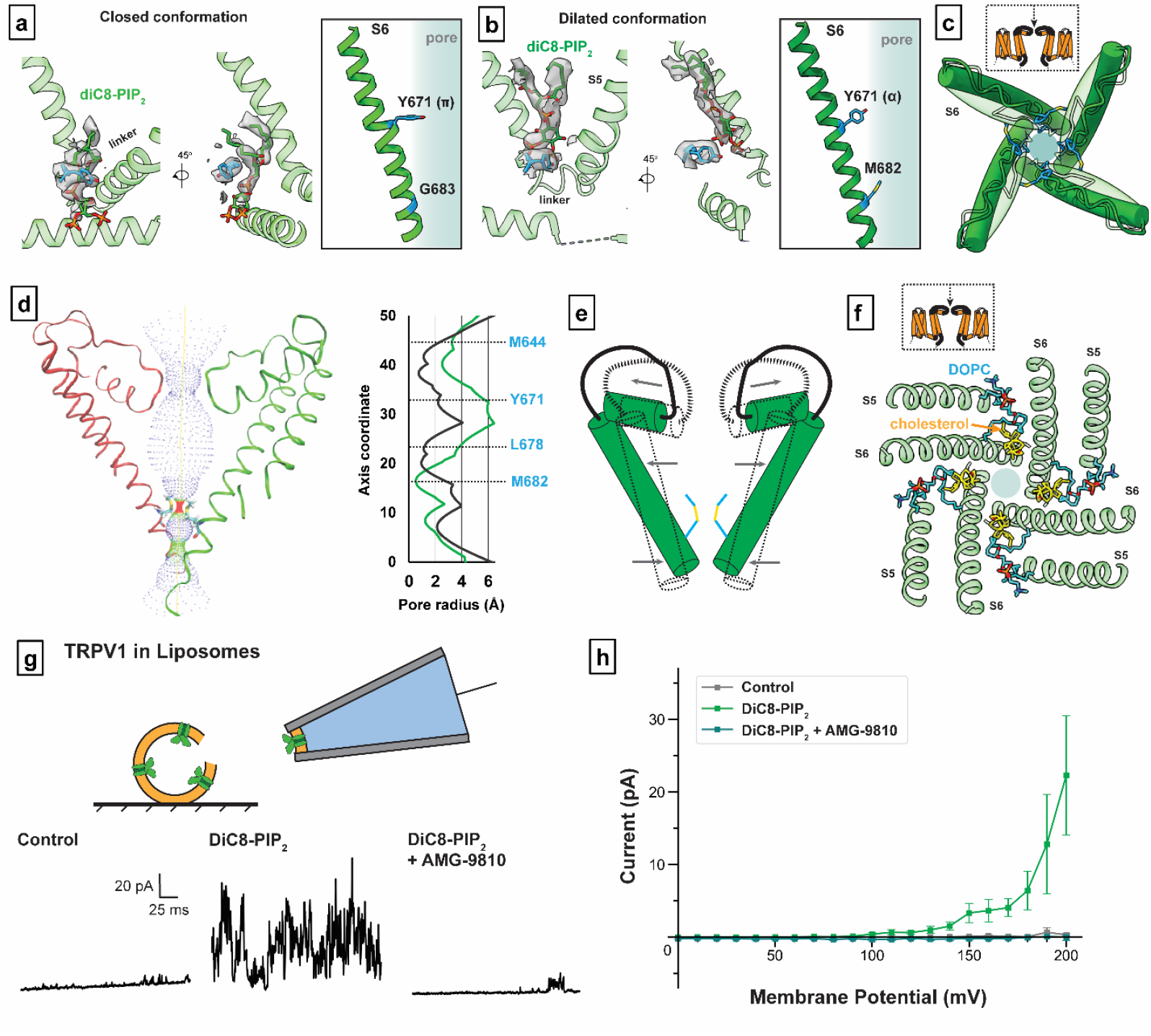
DiC8-PIP_2_ is a partial potentiator of TRPV1 activity. **(a, b)** VBP with diC8-PIP_2_ bound in **(a)** the closed conformation and **(b)** the dilated conformation. **(c)** Top-down view of the TRPV1 pore in the closed conformation (transparent green) and the dilated conformation (dark green). M644 (blue) of the selectivity filter is highlighted. **(d)** (left) Pore profile of TRPV1 in the dilated conformation. (right) Pore radii of closed (black) and dilated (green) states. **(e)** Schematic of the pore movements demonstrating the dilation of the upper portion of the pore and constriction of the lower portion. **(f)** Top-down view of TRPV1 showing the binding of DOPC (blue) and cholesterol (yellow). **(g)** Top: Schematic showing excised inside-out patch clamp recording configuration. Bottom: Sample TRPV1 currents evoked by application of control (bathing) solution, DiC8-PIP_2_, or DiC8-PIP_2_ plus antagonist AMG-9810 (V_h_ = +200 mV). **(h)** Summary current-voltage relationships showing that DiC8-PIP2 reliably activates TRPV1. Data are graphed as mean +/- SEM, n = 10.

In the closed state, channels with diC8-PIP_2_ adopt a conformation like that containing full-length PIP_2_ (Figure 3a), with the acyl chains of the bound lipid preventing reorientation of Y511 or inward movement of the S4-S5 linker. In the dilated state, diC8-PIP_2_ sits higher up in the vanilloid pocket such that the inositol headgroup sits above Y511, which is reorientated toward the pocket (Figure 3b). Moreover, Y671 adopts an α-helical conformation and M682 and is oriented towards the pore axis. These features are hallmarks of a channel in which the resident PI lipid has been displaced (i.e., the pocket is empty or occupied by a vanilloid agonist). However, there are some differences between the diC8-PIP_2_ dilated state and a typical vanilloid agonist-bound structure. Most notably, in the diC8-PIP_2_ dilated state the angle of the S6 helix is more dramatically tilted such that the voltage sensor-like domain (S1-S4) is pivoted outward away from the central axis, and the upper region of the ion permeation pathway, including the selectivity filter, is expanded and stabilized by cholesterol and another membrane lipid (likely DOPC) (Figure 3e and 3f, Supplementary Figure 4). At the same time, pivoting of the S6 helix makes the new restriction site formed by M682 narrower. This dilated structure is consistent with the functional effects of diC8-PIP_2_, which enhances capsaicin sensitivity, but by itself activates the channel only at highly depolarized states (e.g., +150 mV membrane potential)^8,9^ (Figure 3g-3i).

### Channel activation by LPA

LPA is an endogenous bioactive lipid that mediates a host of physiologic responses, including cellular migration and proliferation, inflammation, and pain^14^. While LPA is known to activate metabotropic receptors (GPCRs), it has also been shown to activate or potentiate ionotropic receptors, including TRPV1^10,14,15^. Indeed, LPA activates recombinant TRPV1 channels in reconstituted liposomes^7^ and elicits TRPV1-dependent pain-related behavior in mice^10^. Thus, we are interested in determining how this endogenous pro-algesic lipid binds to TRPV1, and whether and how this involves interaction with the vanilloid site.

To visualize the lipid-channel complex, we added LPA to nanodisc-reconstituted TRPV1 and determined a consensus cryo-EM map to a resolution of 3.0 Å. Through symmetry expansion and focused classification, we captured LPA bound within the vanilloid pocket in multiple stoichiometries, ranging from apo to all 4 subunits occupied (Figures 4 and 5, Supplementary Figure 5). LPA displaces the endogenous PI lipid by occupying the upper region of the pocket that binds aliphatic tails (Figure 4a). As in the case of vanilloid agonists, Y511 flips toward the VBP so that its hydroxyl group forms an H-bond with the ester carbonyl of the LPA tail (Figure 4c). Moreover, S512, T550, and Y554 engage in H-bond interactions with the phosphate moiety of LPA, while the free hydroxyl of the glycerol backbone interacts with N551. Further comparison between bound LPA and RTX shows that both form H-bonds with Y511 and T550 (Figure 4c). An important difference is that the 4-OH of the RTX vanilloid moiety engages in a bridging H-bond network between R557 and E570, two residues that form a salt bridge when TRPV1 is fully opened^5^. This reflects the fact that the RTX 4-OH sits lower into the pocket, closer to R557 and E570, compared to the LPA headgroup. Therefore, the RTX headgroup may be better suited for priming this salt bridge interaction as compared to LPA, which may help to explain the greater efficacy of RTX as an agonist. Together, these structures demonstrate that LPA utilizes similar molecular interactions as vanilloid agonists, such as RTX or capsaicin, for binding to TRPV1 and initiating gating. In the LPA-bound structure, the selectivity filter remains unchanged from the resting state, but we do see that S6 adopts an α helical configuration, resulting in rotation of the lower half of the S6 helix by one residue (Figure 4b), consistent with LPA acting as an agonist.

**Figure 4.**
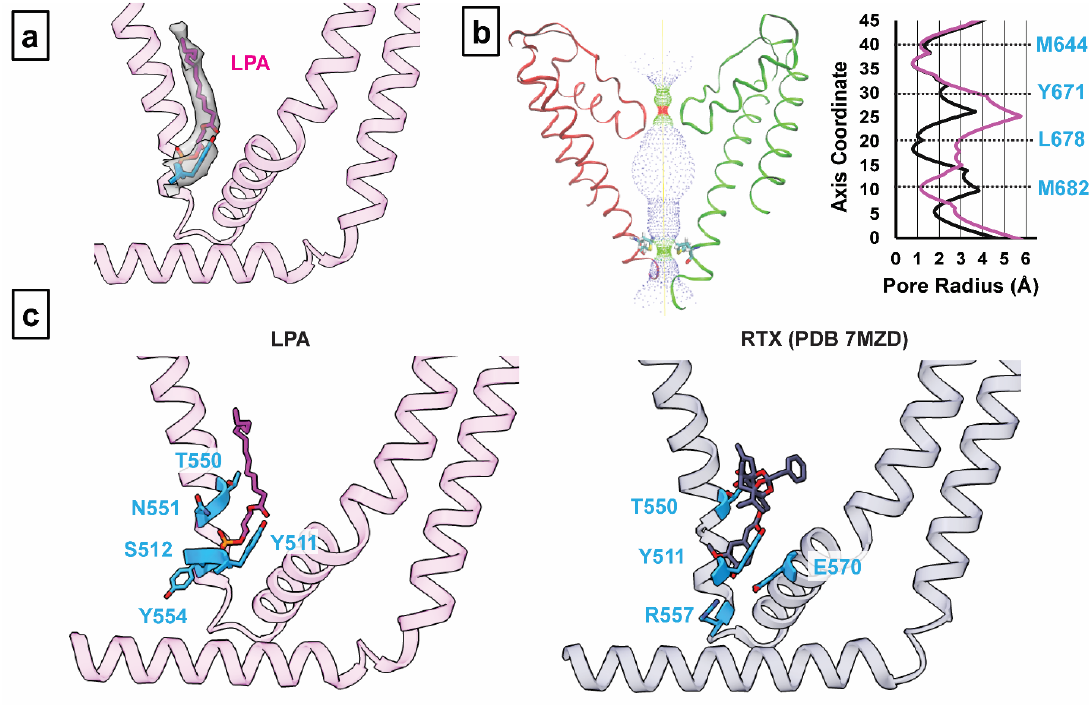
LPA binding to TRPV1. **(a)** VBP with LPA bound. **(b)** (left) Pore profile of TRPV1 with LPA bound. (right) Pore radii show profiles for LPA (magenta) compared to the corresponding apo state (black). **(c)** Molecular interactions between ligand headgroups and TRPV1. Residues that form electrostatic interactions with ligand functional groups (within 3.5 Å) for LPA and RTX (PDB 7MZD) are shown as blue.

**Figure 5.**
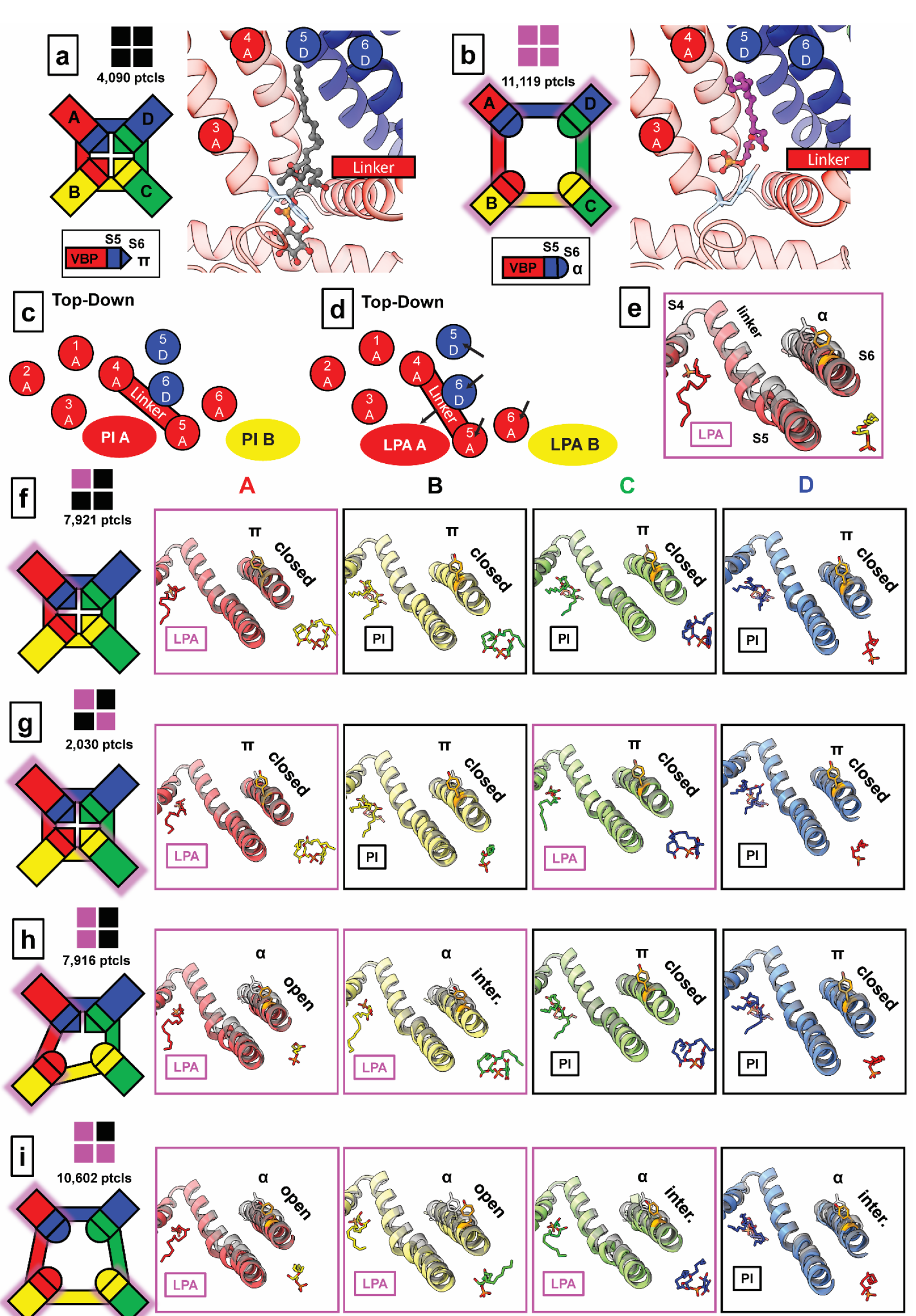
Sub-stoichiometric states of LPA binding. **(a)** The closed configuration of TRPV1 with all four subunits occupied by PI lipid. **(b)** The open configuration TRPV1 with LPA bound to all four subunits. The VBPs of the A monomers are shown to demonstrate the domain-swap architecture that comprises the VBP (S3, S4, and S4-S5 from A; S5 and S6 from D). **(c, d)** Schematic of the binding pocket from a top-down view highlighting the domain-swap architecture and helical movements between the (c) closed and (d) open conformations. **(e)** Structural changes associated with the binding of LPA compared to apo (transparent black). **(f-i)** Sub-stochiometric states of LPA binding. Leftmost panels show a cartoon representation of the TRPV1 tetramer indicating the functional state of each VBP monomer. Monomers are labeled anticlockwise; LPA-occupied monomers are shadowed with magenta. Functional state of the pore is indicated as “closed” (π helix at Y671), “open” (α helix at Y671 with S6 moved away from the pore to the same extent as fully occupied LPA), and “intermediate” (“inter.”, α helix at Y671 with S6 positioned between that of apo and fully occupied LPA).

### LPA substates reveal allosteric gating

In addition to the two structures described above, we also captured structures of TRPV1 with sub-stoichiometric LPA binding, including 1 LPA, 2 LPAs bound to either neighboring or opposite subunits, or 3 LPAs (Figure 5, Supplementary Figure 5). With resting and fully liganded structures representing closed and open configurations of the pore (S6) helices, respectively (Figure 5a and 5b), we examined intermediate conformational states associated with sub-stochiometric LPA binding (the four subunits are denoted as A through D in a counterclockwise orientation as viewed from the extracellular face).

Binding of one LPA to subunit A does not induce an appreciable conformational change to TRPV1 other than replacement of the resident PI lipid and flipping of Y511, with no change to the pore configuration (Figure 5f, Supplementary Figure 6a). The same is true when a second LPA binds to the opposite subunit (A and C) (Figure 5g, Supplementary Figure 6b). However, when the second LPA binds to an adjacent subunit (A and B), we see that gating transitions occur in the S6 helices owing to ejection of the resident PI lipid from both neighboring pockets, which communicate by virtue of domain-swap architecture (Figure 5h, Supplementary Figure 6c). Specifically, we see that one S6 helix (S6-A) is in the open configuration and the neighboring helix (S6-B) is in an intermediate position between open and closed because the resident PI lipid in the C subunit prevents full backward movement of S6-B (Supplementary Figure 7). S6-C and S6-D remain in a closed configuration (Figure 5h). The binding of the third LPA induces all four S6 helices to adopt the π-α transition, with the S6-B and S6-C in the open configuration and the S6-D and S6-A helices in intermediate positions (Figure 5i, Supplementary Figure 6d). Together, these observations are consistent with a model in which ligand binding induces allosteric gating movements in the adjacent VBP through domain-swap interactions.

How does binding of LPA promote movement of S6? Reorientation of Y511 toward the VBP is a key step, allowing L674 on the S4-S5 linker to partially occupy the now vacant space, thereby facilitating movement of the S4-S5 linker towards the VBP (Supplementary Figure 6c). Movement of the S4-S5 linker enables the S5 and S6 helices to move away from the pore axis while providing space to accommodate a π-α helix transition in S6, which is associated with the loss of hydrophobic interactions between M682 and the S4-S5 linker (Supplementary Figure 6b). At the same time, reorientation of Y511 is necessary, but not sufficient, to facilitate gating movements, consistent with the observation that TRPV1 antagonists also induce similar reorientation of Y511^5,11^.

While the fully liganded structures obtained with LPA or RTX are similar, the intermediate configurations associated with sub-stoichiometric ligand binding are noticeably different. For example, movements of the S4-S5 linker are smaller when RTX is the agonist, and the S6 helix does not move until all four subunits are occupied^6^. One caveat to comparing LPA and RTX substates is that those observed with RTX were obtained under low-sodium conditions. Indeed, LPA-bound sub-states may be more physiologically relevant because they were obtained under normal ionic conditions, and LPA is an endogenous, physiologically relevant ligand.

From this analysis, we infer that the first LPA binds to TRPV1 randomly, but such binding increases the affinity for binding to the adjacent versus the opposite monomer. This behavior is supported by the observation that the particle numbers of two LPA molecules bound to opposite versus neighboring subunits is about 1:4, consistent with increased propensity for binding to an adjacent monomer.

## Concluding Remarks

Bioactive lipids are believed to play important roles as pro-algesic agents that contribute to inflammatory pain, in part by enhancing TRPV1 function. However, their site(s) of action and relationship to gating has not been directly visualized. Our structures now show that LPA, an important inflammatory lipid, binds to the VBP and displaces the resident PI lipid, further validating this site as a key regulatory locus for the action of both endogenous and exogenous lipophilic agents. While the VBP can accommodate structurally diverse ligands, including agonists and antagonists, their precise location dictates the functional outcome, as exemplified by our analysis of diC8-PIP_2_. In this case, one ligand can bind to the same pocket but with its inositol headgroup located in two distinct positions associated with closed or dilated pore states. This detailed structural insight resolves a prior conundrum by explaining how soluble diC8-PIP_2_ and full-length PIP_2_ can mediate different effects on TRPV1 function. This, together with our observation that the ‘empty pocket’ channel is in an agonist-like state, substantiates the model in which the resident PI lipid stabilizes the closed state, and its removal favors the open state. Importantly, we also find that the ‘empty pocket’ channel is open at temperatures well below its normal thermal activation threshold, supporting the idea that ejection of the resident lipid is also a critical step in heat-evoked TRPV1 activation^5,16^. This is also consistent with a more recent observation that the heat-activated TRPV3 channel lacks a resident lipid in a region analogous to that of the VBP in TRPV1^17^.

Previous electrophysiological studies showed that the charge-reversal mutation, K710D, abrogates LPA-evoked responses, leading to the conclusion that K710, a residue located on the periphery of TRPV1 at the membrane-protein interface, is key in mediating the binding of LPA^10^. However, our cryo-EM data places the binding of LPA in the VBP, about 25 Å away from K710. We propose that K710 is part of a basic tunnel through which LPA accesses the VBP. Indeed, when the TRPV1 structure is subjected to CAVER analysis, a tunnel is seen at the surface of the inner membrane that connects K710 to the VBP (Supplementary Figure 8) and which is composed of several basic residues. The notion that this tunnel represents a conduit for LPA is consistent with the observation that channel activation is most robust when LPA is applied to excised inside-out membrane patches, providing direct access to the intracellular face of the membrane^10^. Thus, K710 may serve as a guiding residue directing LPA into this access tunnel by interacting with the negative charge of the phosphate headgroup on LPA.

More generally, we and others have shown that lipid regulation is a common feature of many TRP channel family members^3^. While the interaction of phosphoinositide lipids with the TRPV1 VBP is perhaps the best described, structural and functional studies suggest that other binding sites may exist. One such example is a proposed interaction between PIP_2_ and positively charged residues in the cytoplasmic carboxy-terminal tail^18^; another is a putative site between neighboring subunits of TRPM8, where PIP_2_ binding may enhance channel activity^19^. The structural details and consequences of phosphoinositide binding to these or other sites remain unresolved.

## Materials and Methods

### Materials

Reagents were purchased from Sigma-Aldrich (St. Louis, MO, USA) unless noted below. 18:1 Lysophosphatidic acid [(2-hydroxy-3-phosphonooxypropyl) (*Z*)-octadeca-9-enoate] was purchased from Cayman Chemical Company (Ann Arbor, MI, USA) for cryo-EM sample prep and diluted in DMSO. DiC8-PIP_2_ (1,2-dioctanoyl-sn-glycero-3-phospho-(1’-myo-inositol-4’,5’-bisphosphate)) was purchased from Avanti Polar Lipids and diluted in aqueous buffer prior to cryo-EM or electrophysiology. Soy polar lipid extract, di18:1 PI, and di18:1 PI(4,5)P_2_, 1,2-dioleoyl-sn-glycero-3-phosphocholine (DOPC) and 1-palmitoyl-2-oleoyl-sn-glycero-3-phospho-(1’-rac-glycerol) (POPG) were also purchased from Avanti Polar Lipids Bio-beads SM2 was purchased from Bio-Rad (Hercules, CA, USA). Freestyle 293 Expression Medium and Expi 293 Expression Medium were purchased from Gibco, along with Expi293F cells and the ExpiFectamine 293 Transfection Kit. Sf9 and insect cell culture media were purchased from Expression Systems (Davis, CA, USA). Fetal bovine serum was purchased from PEAK (Wellington, CO, USA) and bovine calf serum was purchased from HyClone (Marlborough, MA, USA). HEK293 GnTI^-^ cells and HEK293T cells were purchased from ATCC (Manassas, VA, USA). DH5α competent cells were purchased from New England Biolabs (Ipswich, MA, USA). Quantifoil® R1.2/1.3 Au 300 mesh grids were purchased from Quantifoil Micro Tools GmbH (Großlöbichau, Germany).

### Brominated lipid synthesis

3-((9,10-dibromooctadecanoyl)oxy)-2-(((9,10-dibromooctadecanoyl)oxy)methyl)propyl ((1S,2R,3R,4S,5S,6R)-2,3,4,5,6-pentahydroxycyclohexyl) phosphate (**PI-Br**_**4**_) and (1R,2R,3S,4R,5R,6S)-4-(((3-((9,10-dibromooctadecanoyl)oxy)-2-(((9,10-dibromooctadecanoyl)oxy)methyl)propoxy)oxidophosphoryl)oxy)-3,5,6-trihydroxycyclohexane-1,2-diyl bis(phosphate) (**PIP**_**2**_**-Br**_**4**_) were synthesized from PI and PIP_2_, respectively using Br_2_ as previously described^12^. 0.5 mg di18:1-PI(4,5)P_2_ or di18:1-PI was dissolved in 0.5 mL CHCl_3_ and stirred on ice in a glass vial. Br_2_ (stoichiometric with the number of double bonds in the lipid) was added to the vial with a glass syringe. The vial was flushed with argon and sealed. The reaction was allowed to continue in the dark with stirring for 1 hr. Solvent and any excess bromine was removed by application of vacuum in the dark overnight. Brominated lipids were stored at -80 ºC until use.

### Protein purification and nanodisc reconstitution

Recombinant minimal functional rat TRPV1 (residues 110-603 and 627-76) as expressed in HEK293 GnTI^-^ cells and purified as previously described^20^. The membrane scaffold protein, MSP2N2, for nanodisc reconstitution was expressed in *E. coli* as previously described^5^.

Nanodisc reconstitution of purified minimal TRPV1 with soybean polar lipids or a defined lipid composition (purchased from Avanti Polar Lipids), was performed following the protocol described previously^5^. Samples were prepared using the following modifications from these published protocols:

#### Empty pocket

5 μM capsaicin in column buffer was applied to the TRPV1-MBP-bound amylose column for 30 min. Column was washed 10 times with 3 × column volume. Protein was eluted and then reconstituted into defined-lipid nanodiscs containing (by mol%) 8% cholesterol, 36.8% 1-palmitoyl-2-oleoyl-phosphatidylglycerol (POPG), and 55.2% dioleoyl-phosphatidylcholine (DOPC). Samples were immediately taken for cryo-EM grid preparation after size-exclusion chromatography.

#### Brominated phosphoinositides

Samples were prepared like the ‘empty pocket’ prep. Nanodiscs were prepared using a defined-lipid composition containing 8% cholesterol, 10% brominated phosphoinositide (PI-Br_4_ or PIP_2_-Br_4_), 32.8% POPG, and 49.2% DOPC.

#### DiC8-PIP_2_

Sample was prepared using the ‘empty pocket’ prep. DiC8-PIP_2_ was applied to sample and then to grid as stated below.

#### LPA

Sample was prepared using the standard protocol with nanodiscs containing soybean polar lipids (i.e., resident lipid was not removed).

### Cryo-EM sample preparation and data acquisition

To prepare cryo-EM grids, 3 μL TRPV1-nanodiscs was applied to glow-discharged Quantifoil R1.2/1.3 Au 300 mesh grids covered in holey carbon film (Quantifoil Micro Tools GmbH) and blotted with Whatman 1 filter paper on a Vitrobot Mark IV (FEI company) with a 4.5 s blotting time, 4 blot force, and 100% humidity, and subsequentially plunge-frozen in liquid ethane cooled by liquid nitrogen. Sample was imaged with a Titan Krios microscope (ThermoFisher FEI) operated at 300 kV and equipped with a post-column Bio Quantum energy filter with zero-loss energy selection slit set to 20 eV and a K3 camera (Gatan) either at UCSF or using the Cryo-EM Consortium at Stanford SLAC (S^2^C^2^) as stated below. Data collection was carried out with SerialEM software^21^. The detailed collecting parameters, including dose rate, total dose, and total frames per movie stack, etc. are summarized in Supplementary Tables 1-4. Specific conditions not stated above for each of the samples are described below.

#### Empty-pocket TRPV1 at 4° C (Grid 1)

Empty-pocket TRPV1-nanodisc (described above) was kept on ice prior to applying to grids in a Vitrobot kept at 4° C during grid preparation. Data was collected at S^2^C^2^ using TEM Gamma.

#### Empty-pocket TRPV1 at 25° C (Grid 2)

Empty-pocket TRPV1-nanodisc was kept on ice, briefly warmed to room temperature prior to applying to grids in a Vitrobot kept at 25° C. Data was collected at S^2^C^2^ using TEM Beta.

#### PI-Br_4_ (Grid 3) and PIP_2_-Br_4_ (Grid 4)

TRPV1-nanodisc containing brominate phosphoinositide was kept on ice and applied to grids using a Vitrobot kept at 25° C. Data collected at UCSF.

#### DiC8-PIP_2_ (Grid 5)

DiC8-PIP_2_ was dissolved in buffer containing 20 mM HEPES (pH 7.5), 150 mM NaCl, and 0.1 mM TCEP*HCl to a stock concentration of 1 mM. DiC8-PIP_2_ was applied to TRPV1-nanodiscs to a final concentration of 50 μM for 30 min on ice before applying sample to grids using a Vitrobot kept at 15° C. Data collected at UCSF.

#### LPA (Grid 6)

LPA was dissolved in DMSO to a working stock of 5 mM. LPA was applied to a final concentration of 50 μM (1% vehicle) to TRPV1-nanodiscs kept at room temperature for 30 min. Sample was applied to grids using a Vitrobot kept at 25° C. Data collected at UCSF.

### Image processing

Cryo-EM data processing is illustrated in Supplementary Figures 9-14 and dFSC curves are provided in Supplementary Figure 15. In general, motion correction on movie stacks was processed on-the-fly using MotionCorr2^22^ and binned 2 × 2 with Fourier cropping to 0.835 Å/pixel (UCSF Krios) or to 0.68 Å/pixel (S^2^C^2^). Dose-weighted micrographs were visually inspected to remove bad micrographs before further processing by cryoSPARC^23^. Patch-based CTF Estimation was performed in cryoSPARC. Micrographs with estimated resolution poorer than 4.5 Å were discarded. EMD-8118 (apo TRPV1 in nanodisc) was used to make 25 templates for template picking. Picks were extracted and binned 4 × 4 by Fourier cropping and reference-free 2D classification was used to remove non-TRPV1-nanodisc picks. Extracted particles were then subjected to reference-based 3D classification (ref. EMD-8118, low-pass filtered 12 Å) RELION^24^ to remove low-resolution and featureless particles. Further processing for each dataset continued as stated below. Resolutions were determined according to the gold standard Fourier Shell Correlation = 0.143 criterion^25^.

#### ‘Empty pocket’ TRPV1 4° C

Particles were enriched for high-resolution features using symmetry expansion followed by focused classification as previously described^26^. Specifically, particles were refined using 3D Auto Refine with C4 symmetry in RELION using EMD-8118 as a reference (low-pass filter 12 Å). These refined particles were symmetry-expanded using C4 symmetry and a mask focused on the vanilloid binding pocket was used for background subtraction, followed by 3D classification on the symmetry-expanded particles. Classification parameters: ref. EMD-8118 (no low-pass filter), Regularization Parameter T = 80, symmetry C1, no image alignment. Particles without defined vanilloid pocket features were discarded. Particles containing monomers with 4 distinguishable pockets were than taken to cryoSPARC for non-uniform refinement (C4 symmetry) and then sharpened in PHENIX^27^ using half-map sharpening. Final resolution: 2.9Å.

#### ‘Empty pocket’ TRPV1 25° C

Selected particles from 3D classification were refined in cryoSPARC using non-uniform refinement and C4 symmetry. Final resolution: 3.7 Å.

#### PI-Br_4_

Selected particles from 2D classification were refined in RELION using C4 symmetry (reference EMD-8118, low-pass filtered 12 Å) to 2.5 Å. Particles were then 3D classified in RELION using no image alignment and selected 3D classes with well-defined lipid features were refined in RELION as before. To further clarify bromine features, particles were subjected to the symmetry expansion and focused classification regiment as stated for ‘Empty Pocket’ TRPV1 at 4° C. Classification parameters: regularization parameter T = 80, no low-pass on reference, no image alignment. Classes with all 4 monomers containing well-resolved phosphoinositide densities were refined in cryoSPARC using non-uniform refinement and C4 symmetry. The consensus map reached 2.2 Å resolution.

The consensus map showed a density projecting from the main-chain acyl density indicative of a secondary conformation for the lipid tail. Phosphoinositide tail conformations were further resolved with another round of refinement in RELION and symmetry expansion and focused classification (regularization parameter T = 80, no low-pass on reference, Arnold, W. R., et al., 18 September, 2023 – preprint copy – BioRxiv no image alignment). Two classes emerged with distinct tail conformations, named Conformation 1 (tail projecting upward in the binding pocket as seen in other structures) and Conformation 2 (tail projecting towards the membrane). Conformation 1 particles were combined, and Conformation 2 particles were combined, and the subtracted particles (focused on the vanilloid pocket) were refined in RELION using EMD-8118 as the reference and a low pass filter of 12 Å and sharpened using Post Process in RELION. Final resolution reached 2.3 Å for both conformations. Conformation 1 is used in the main text figures as a representative of PIBr_4_.

#### PIP_2_-Br_4_

Particles selected from 2D classes were extracted and then refined using cryoSPARC reference-based (EMD-8118) non-uniform refinement. As the sample is very homogenous, 3D classification did not improve data quality, and so was not used in the final data. Map resolved to 2.4 Å.

#### DiC8-PIP_2_

Particles were refined in RELION using EMD-8118 as a reference low-pass filtered to 12 Å. Particles were then subjected to the symmetry expansion and focused classification regiment as stated above (regularization parameter T = 40, reference low-pass filtered to 12 Å, no image alignment). Low-resolution particles were excluded and two conformations of diC8-PIP_2_ emerged, a closed conformation and a dilated-state conformation. Particles containing all four monomers containing either diC8-PIP_2_ in the closed state or in the dilated state were taken further for refinement in RELION (reference EMD-8118, low-pass filtered 12 Å) and then sharpened using Post Process in RELION. Final resolution for closed was 3.0 Å and for dilated 3.6 Å.

#### LPA

The consensus map for LPA resolved to 3.0 Å. To further sperate LPA-bound from PI-bound subunits, all particles in the final dataset were subjected to symmetry expansion followed by focused classification as stated above. Classification parameters: ref. EMD-8118 (no low-pass filter), Regularization Parameter T = 80, symmetry C1, no image alignment. Classes were whitelisted for further processing based on clarity of the ligand features, and the ligand content (LPA or lipid) was visually assigned to each class. Particles were grouped based on the number of LPA bound (0, 1, 2, 3, or 4) and the arrangement (neighboring versus opposite pockets) of how the ligand is bound in tetrameric particles. New star files were generated for each group to calculate a 3D reconstruction followed by further refinement.

Particles with either 0 or 4 LPA molecules bound were refined in cryoSPARC using non-uniform refinement with C4 symmetry, followed by PHENIX^27^ half-map sharpening. For other ligand-bound particles, transmembrane-region focused and local refinement was performed on the pre-aligned particles in RELION. The resulting maps were sharpened using RELION Post-Process and the transmembrane mask was used as the solvent mask. Reconstructions with LPA bound in one, two neighboring, or three subunits were refined without symmetry. For reconstruction with 2 LPA bound in two opposite subunits, particles were first subjected to 3D classification using 1 class, Regularization Parameter T = 40, symmetry C2, transmembrane mask, and local searches to obtain the initial angles. The star file and resulting map were used in 3D Auto Refine as the input star file and the reference (low-pass filtered to the classification resolution of 4.1 Å), respectively, and refinement was carried out using a transmembrane mask and local searches. The resulting map was sharpened using RELION Post-Process and the transmembrane mask as the solvent mask.

## Model building

Resting TRPV1 (PDB-7L2P) was used as the starting model and docked into the sharpened maps using UCSF Chimera^28^, followed by manual adjustment based on the resolvable features of the maps. PDB and molecular restraint files for ligands were generated in PHENIX using eLBOW, and then manually docked into the ligand densities. For the general ‘resident lipid’, di-palmitoyl phosphatidylinositol was used as the starting structure and then the tails were shortened to fit the resolvable density. Models were refined using several iterations of PHENIX Real Space Refine and manual adjustments in COOT^29^. The quality of the refined models was determined using the wwPDB validation server^30^ and the results are shown in Supplementary Tables 1-4.

## Difference map analysis

Difference map analysis was used to emphasize densities for bromine atoms and phosphate groups in the PI-Br_4_ (Conformation 1) and PIP_2_-Br_4_ maps. The final PDB models were used as the basis to generate calculated maps using the MolMap function in Chimera^28^ at 2.3 Å resolution for PI-Br_4_ and 2.4 Å for PIP_2_-Br_4_ (Supplemental Figure 10). As resolution and B factors vary throughout the lipid densities, analyses were performed on a focused area around the bromine atoms (PI-Br_4_ and PIP_2_-Br_4_) or the inositol headgroup (for PIP_2_-Br_4_). For PI-Br_4_, the calculated map was generated from a PDB file that contained carbons 8-11 of the acyl tail and lacking the bromine atoms. For PIP_2_-Br_4_, a lipidic density comes near to bromine atoms of PIP_2_-Br_4_ and forms a connecting density. Therefore, the analysis PDB contained carbons 8-11 of the PIP_2_-Br_4_ acyl tail (lacking bromine) and the immediate carbon density of the other lipid. The corresponding densities in the real maps were then isolated and the TEMPy:DiffMap function in the CCPEM^31,32^ suite was used to generate difference maps between the real densities and the calculated densities.

## Pore radius determination, tunnel analysis, electrostatics calculation, and structural figures

The pore radii were determined using the HOLE program^33^ and plotted in Excel. Tunnels within TRPV1 were identified, visualized, and made into a figure using the program CAVER^34^. Surface electrostatic charge determination was performed, visualized, and made into a figure in PyMOL. All other structural figures were made using UCSF ChimeraX^35,36^ and Adobe Illustrator.

## TRPV1 proteoliposome preparation

Liposomes were prepared as reported previously^7^. Minimal functional rat TRPV1 with an N-terminal 8xHis-MBP tag was expressed in Expi-293F cells for two days using the Expifectamine-293 Transfection Kit (Thermo Fisher Scientific). Transfected cells were then harvested by centrifugation at 3,000xg for 10 minutes at 4°C, with the supernatant decanted and cell pellets flash-frozen in liquid nitrogen and stored at - 80°C until use. To purify TRPV1, pellets were thawed and resuspended with buffer containing 200 mM NaCl, 50 mM HEPES pH = 8, 2 mM TCEP, 10% glycerol, and protease inhibitors (Pierce tablet). 20 mM DDM (Anatrace) was added to extract TRPV1, while incubating on a rotator for 2 hours at 4°C. Samples were spun at 20,000xg for 1 hour at 4°C, with the supernatant being collected, filtered at 0.2 μm, and combined with ∼1 mL amylose resin (New England BioLabs) for at least 1 hour of affinity binding. Beads were poured over a Poly-Prep column (Bio-Rad) and washed with ∼20 mL purification buffer (containing 200 mM NaCl, 50mM HEPES pH = 8, 2 mM TCEP, 10% glycerol, 1 mM DDM, and 10 μg/mL defined lipid mixture i.e. DOPC:POPG:Cholesterol as done with nanodiscs) to remove impurities. TRPV1 was eluted with purification buffer plus 20 mM maltose.

Additional defined lipid mixture was dried down under nitrogen gas and stored in a vacuum desiccator one day prior to the liposome prep. The dried lipid was dissolved in buffer containing 200 mM NaCl, 5 mM MOPS pH = 7, and 2 mM TCEP, to achieve a final concentration of 5 mg/mL. The lipid stock was left to sit for 30 minutes, sonicated for 10 minutes, and subjected to ten freeze-thaw cycles with liquid nitrogen and hot water to ensure lipid dispersion. Lipids were further destabilized by addition of 4 mM DDM and left to rotate for 30 minutes at room temperature. The resulting lipid-detergent stock was combined with eluted TRPV1 to achieve the desired 1:5 protein-to-lipid mass ratio and left to equilibrate for 1 hour at room temperature on a rotator. Bio-Bead SM-2 resin was then added in four doses (60 mg, 60 mg, 150 mg, 300 mg per 10 mg lipid sample) with 1 hour room-temperature rotator incubations in-between. After the last Bio-Bead incubation, the mixture was left overnight (over 15 hours) and transferred to a 4°C rotator. The next day, Bio-Beads were removed by a Poly-Prep column (Bio-Rad) and washed with a small volume of minimal buffer (containing 200 mM NaCl and 5 mM MOPS pH = 7). Liposomes were pelleted at 100,000xg for 1 hour at 4°C. Liposome pellets were resuspended in 200 uL of minimal buffer, divided into 13 uL aliquots, flash-frozen in liquid nitrogen, and stored at -80°C until use.

## Liposome Electrophysiology

Liposome electrophysiology was performed as previously described^7^. The day before a patch clamp session, one aliquot of frozen liposomes was thawed, supplemented with an equal volume of minimal buffer (containing 200 mM NaCl and 5 mM MOPS pH = 7) plus 40 mM sucrose, plated on a glass coverslip, and dehydrated in a vacuum desiccator for at least 50 minutes. 100 μL of minimal buffer were added to the dried liposomes to rehydrate them overnight. Prior to patching, 5 uL of rehydrated liposomes were pipetted onto additional coverslips with 100 uL of minimal buffer on them and left still for two hours to allow liposomes to adhere to the glass.

Bathing solution was delivered by gravity perfusion and contained 140 mM NaCl, 5 mM KCl, 20 mM HEPES pH = 7.6 and 2 mM MgCl2. Internal solution was kept identical to bathing solution. Pipette tips were pulled from BF150-86-10 borosilicate capillaries (Sutter Instruments) using a P-97 micropipette puller (Sutter Instruments) and fire-polished using a MF-830 microforge (Narishige), keeping tip resistances between 3 and 10 MΩ. Coverslips with liposomes were transferred to an IX71 inverted microscope setup (Olympus). The electrode tip was pressed up against a liposome using an MP-285 micromanipulator (Sutter Instruments) and pressure was applied orally until a giga-Ohm seal was achieved. The tip was then slowly retracted until just a patch of membrane was retained, accessing the inside-out patch configuration. The patch was then placed in front of the SmartSquirt microperfusion system (AutoMate Scientific) to apply chemical ligands. Voltage steps between -200 mV and +200 mV were applied in 10 mV increments, and currents were recorded, via an AxoPatch 200B amplifier (Molecular Devices) and a Digidata 1550B digitizer (Molecular Devices). Signals were acquired at 20 kHz and filtered at 5 kHz. Data were analyzed post-hoc in pClamp (Molecular Devices), Excel (Microsoft), and Python. TRPV1-specific currents were calculated by subtracting the leak of the patch, estimated as the lowest magnitude peak in an all-point histogram of current amplitude in a given potential, from the mean current measured over the given potential.

## End Matter

### Author contributions

WRA conceived the project, performed cryo-EM experiments, and analyzed data. ASM performed and analyzed the electrophysiology experiments. FM synthesized the brominated phosphoinositide lipids under the guidance of AF. YC and DJ provided advice and guidance throughout. All authors contributed to manuscript preparation.

## Acknowledgements

We thank Dr. David Bulkley and Glenn Gilbert of the UCSF EM Core Facility for assistance with cryo-EM data acquisition, and Ms. Yifei Chen for help with data processing, and Dr. T. Rosenbaum for advice on LPA pharmacology. Parts of the cryo-EM data were collected at the Stanford-SLAC Cryo-EM Center (S^2^C^2^) supported by the NIH Common Fund Transformative High Resolution Cryo-Electron Microscopy program (U24 GM129541. This work was supported by an IRACDA NIH Training Award (K12GM081266 to W.R.A.) and grants from the NIH (R35NS105038 to D.J., 1R35GM140847 to Y.C., and P50 GM082545 and 1DP2-GM110772 to A.F.). Y.C. is an investigator of the Howard Hughes Medical Institute. A.S.M. is supported by a Postgraduate Scholarship – Doctoral award, provided by the Natural Sciences and Engineering Research Council of Canada (NSERC). A.F. was supported by a Faculty Scholar grant from the HHMI and is currently a Chan Zuckerberg Biohub investigator. F.R.M. was supported by a postdoctoral fellowship from The Jane Coffin Childs Memorial Fund for Medical Research. Structural biology applications used in this project were compiled and configured by SBGrid.

## Data availability

PDB structures EMD maps be accessed by the following ascension codes, respectively. Empty pocket 4°C (8U3J, 41864), empty pocket 25°C (8U3L, 41866), PI-Br_4_ consensus (8U4D, 41879), PI-Br_4_ Conformation 1 (8U3A, 41855), PI-Br_4_ (8U3C, 41857), PIP_2_-Br_4_ (8U43, 41873, diC8-PIP_2_ closed (8U30, 41848), diC8-PIP_2_ dilated (8U2Z, 41847), LPAx0 (8T0E, 40941), LPAx1 (8T0Y, 40949)I, LPAx2 opposite (8T10, 40951), LPAx2 neighboring (8T3L, 41005), LPAx3 (8T3M, 41006), and LPAx4 (8T0C, 40940).

## References

1 Estacion, M., Sinkins, W. G. & Schilling, W. P. Regulation of Drosophila transient receptor potential-like (TrpL) channels by phospholipase C-dependent mechanisms. J Physiol 530, 1–19 (2001). 10.1111/j.1469-7793.2001.0001m.x

2 Huang, J. et al. Activation of TRP channels by protons and phosphoinositide depletion in Drosophila photoreceptors. Curr Biol 20, 189–197 (2010). 10.1016/j.cub.2009.12.019

3 Rohacs, T. Phosphoinositide regulation of TRP channels. Handb Exp Pharmacol 223, 1143–1176 (2014). 10.1007/978-3-319-05161-1_18

4 Chuang, H. H. et al. Bradykinin and nerve growth factor release the capsaicin receptor from PtdIns(4,5)P2-mediated inhibition. Nature 411, 957–962 (2001). 10.1038/35082088

5 Gao, Y., Cao, E., Julius, D. & Cheng, Y. TRPV1 structures in nanodiscs reveal mechanisms of ligand and lipid action. Nature 534, 347–351 (2016). 10.1038/nature17964

6 Zhang, K., Julius, D. & Cheng, Y. Structural snapshots of TRPV1 reveal mechanism of polymodal functionality. Cell (2021). 10.1016/j.cell.2021.08.012

7 Cao, E., Cordero-Morales, J. F., Liu, B., Qin, F. & Julius, D. TRPV1 channels are intrinsically heat sensitive and negatively regulated by phosphoinositide lipids. Neuron 77, 667–679 (2013). 10.1016/j.neuron.2012.12.016

8 Lukacs, V. et al. Dual regulation of TRPV1 by phosphoinositides. J Neurosci 27, 7070–7080 (2007). 10.1523/JNEUROSCI.1866-07.2007

9 Rohacs, T. Phosphoinositide regulation of TRPV1 revisited. Pflugers Arch 467, 1851–1869 (2015). 10.1007/s00424-015-1695-3

10 Nieto-Posadas, A. et al. Lysophosphatidic acid directly activates TRPV1 through a C-terminal binding site (vol 8, pg 78, 2012). Nat Chem Biol 8, 737–737 (2012). 10.1038/nchembio0812-737c

11 Neuberger, A. et al. Human TRPV1 structure and inhibition by the analgesic SB-366791. Nat Commun 14, 2451 (2023). 10.1038/s41467-023-38162-9

12 Moss, F. R., 3rd et al. Brominated lipid probes expose structural asymmetries in constricted membranes. Nat Struct Mol Biol 30, 167–175 (2023). 10.1038/s41594-022-00898-1

13 Ufret-Vincenty, C. A., Klein, R. M., Hua, L., Angueyra, J. & Gordon, S. E. Localization of the PIP2 sensor of TRPV1 ion channels. J Biol Chem 286, 9688–9698 (2011). 10.1074/jbc.M110.192526

14 Geraldo, L. H. M. et al. Role of lysophosphatidic acid and its receptors in health and disease: novel therapeutic strategies. Signal Transduct Target Ther 6, 45 (2021). 10.1038/s41392-020-00367-5

15 Pan, H. L., Zhang, Y. Q. & Zhao, Z. Q. Involvement of lysophosphatidic acid in bone cancer pain by potentiation of TRPV1 via PKCepsilon pathway in dorsal root ganglion neurons. Mol Pain 6, 85 (2010). 10.1186/1744-8069-6-85

16 Cao, E. H., Liao, M. F., Cheng, Y. F. & Julius, D. TRPV1 structures in distinct conformations reveal activation mechanisms. Nature 504, 113-+ (2013). 10.1038/nature12823

17 Singh, A. K. et al. Structural basis of temperature sensation by the TRP channel TRPV3. Nat Struct Mol Biol 26, 994–998 (2019). 10.1038/s41594-019-0318-7

18 Prescott, E. D. & Julius, D. A modular PIP2 binding site as a determinant of capsaicin receptor sensitivity. Science 300, 1284–1288 (2003). 10.1126/science.1083646

19 Yin, Y. et al. Activation mechanism of the mouse cold-sensing TRPM8 channel by cooling agonist and PIP(2). Science 378, eadd1268 (2022). 10.1126/science.add1268

20 Liao, M., Cao, E., Julius, D. & Cheng, Y. Structure of the TRPV1 ion channel determined by electron cryo-microscopy. Nature 504, 107–112 (2013). 10.1038/nature12822

21 Mastronarde, D. N. Automated electron microscope tomography using robust prediction of specimen movements. J Struct Biol 152, 36–51 (2005). 10.1016/j.jsb.2005.07.007

22 Zheng, S. Q. et al. MotionCor2: anisotropic correction of beam-induced motion for improved cryo-electron microscopy. Nat Methods 14, 331–332 (2017). 10.1038/nmeth.4193

23 23 Punjani, A., Rubinstein, J. L., Fleet, D. J. & Brubaker, M. A. cryoSPARC: algorithms for rapid unsupervised cryo-EM structure determination. Nat Methods 14, 290–296 (2017). 10.1038/nmeth.4169

24 Scheres, S. H. RELION: implementation of a Bayesian approach to cryo-EM structure determination. J Struct Biol 180, 519–530 (2012). 10.1016/j.jsb.2012.09.006

25 Rosenthal, P. B. & Henderson, R. Optimal determination of particle orientation, absolute hand, and contrast loss in single-particle electron cryomicroscopy. J Mol Biol 333, 721–745 (2003). doi.org:DOI 10.1016/j.jmb.2003.07.013

26 Arnold, W. R., Asarnow, D. & Cheng, Y. Classifying liganded states in heterogeneous single-particle cryo-EM datasets. Microscopy (Oxf) 71, i23–i29 (2022). 10.1093/jmicro/dfab044

27 Liebschner, D. et al. Macromolecular structure determination using X-rays, neutrons and electrons: recent developments in Phenix. Acta Crystallogr D 75, 861–877 (2019). 10.1107/S2059798319011471

28 Pettersen, E. F. et al. UCSF Chimera--a visualization system for exploratory research and analysis. J Comput Chem 25, 1605–1612 (2004). 10.1002/jcc.20084

29 Emsley, P., Lohkamp, B., Scott, W. G. & Cowtan, K. Features and development of Coot. Acta Crystallographica Section D-Biological Crystallography 66, 486–501 (2010). 10.1107/S0907444910007493

30 Berman, H., Henrick, K. & Nakamura, H. Announcing the worldwide Protein Data Bank. Nat Struct Biol 10, 980 (2003). 10.1038/nsb1203-980

31 Burnley, T., Palmer, C. M. & Winn, M. Recent developments in the CCP-EM software suite. Acta Crystallogr D Struct Biol 73, 469–477 (2017). 10.1107/S2059798317007859

32 Wood, C. et al. Collaborative computational project for electron cryo-microscopy. Acta Crystallogr D Biol Crystallogr 71, 123–126 (2015). 10.1107/S1399004714018070

33 Smart, O. S., Neduvelil, J. G., Wang, X., Wallace, B. A. & Sansom, M. S. HOLE: a program for the analysis of the pore dimensions of ion channel structural models. J Mol Graph 14, 354-360, 376 (1996). 10.1016/s0263-7855(97)00009-x

34 Jurcik, A. et al. CAVER Analyst 2.0: analysis and visualization of channels and tunnels in protein structures and molecular dynamics trajectories. Bioinformatics 34, 3586–3588 (2018). 10.1093/bioinformatics/bty386

35 Goddard, T. D. et al. UCSF ChimeraX: Meeting modern challenges in visualization and analysis. Protein Sci 27, 14–25 (2018). 10.1002/pro.3235

36 Pettersen, E. F. et al. UCSF ChimeraX: Structure visualization for researchers, educators, and developers. Protein Sci 30, 70–82 (2021). 10.1002/pro.3943

