## Supplementary Information for "Structural basis of TRPV1 modulation by endogenous bioactive lipids"

**William R. Arnold1, Adamo Mancino, Frank R. Moss III1,4, Adam Frost1,4,5, David Julius2*, and Yifan Cheng1,3***

[1]Department of Biochemistry and Biophysics, University of California San Francisco, 600 16th St. San Francisco, CA, 94158
[2]Department of Physiology, University of California San Francisco, 600 16th St. San Francisco, CA, 94158
[3]Howard Hughes Medical Institute, University of California San Francisco, San Francisco, CA 94158
[4]Present address: Chan Zuckerberg Biohub, San Francisco, CA 94158
[5]Present address: Altos Labs, Redwood City, CA 94065
*corresponding authors

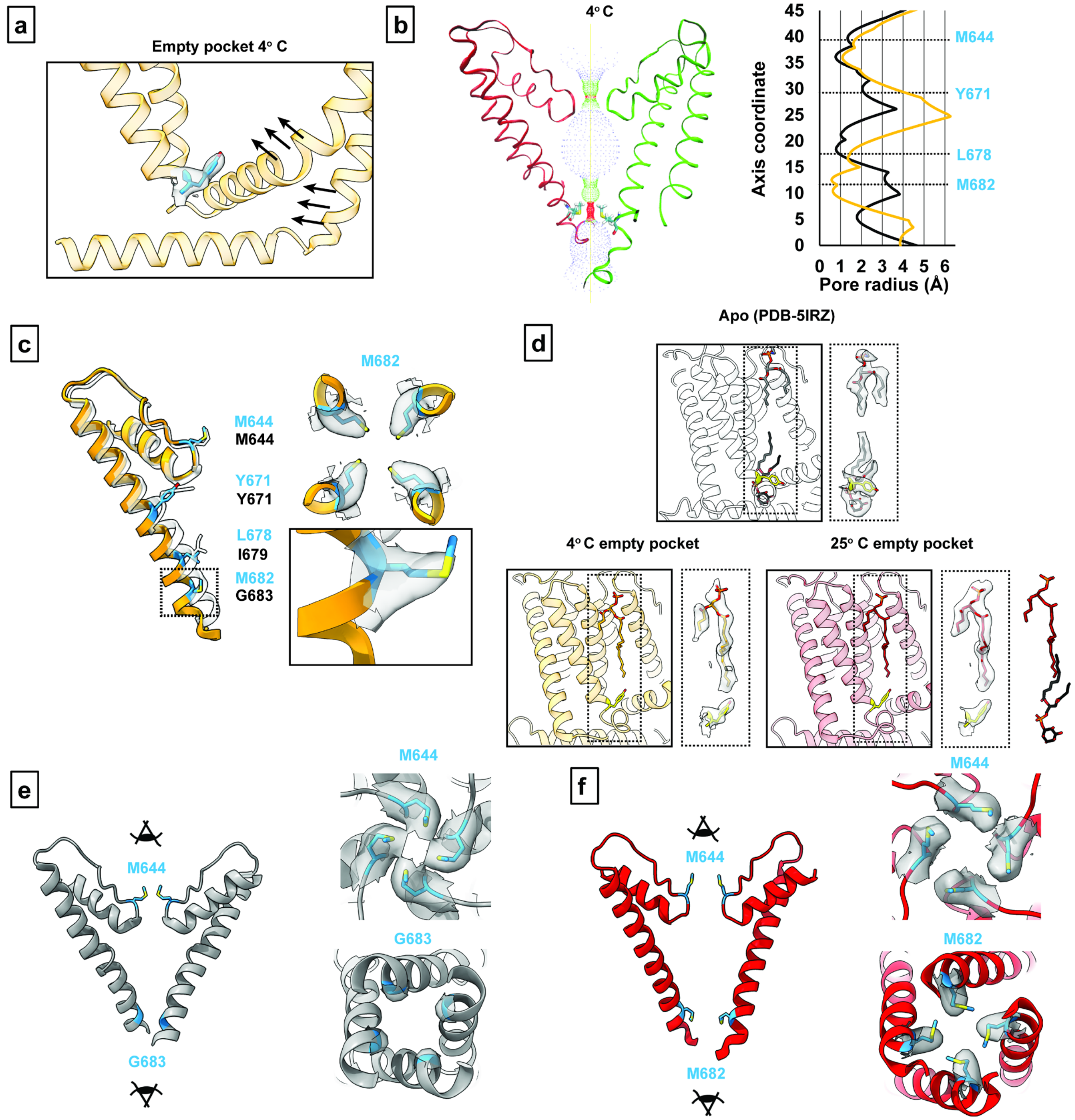


Supplementary Figure 1. *Supplementary TRPV1 empty pocket data*. **(a)** Vanilloid pocket with empty-pocket TRPV1 at 4° C. **(b)** Pore profile for empty-pocket TRPV1 at 4° C. Pore radius was determined using the HOLE program. Black is apo TRPV1 and orange is empty-pocket TRPV1. **(c)** Key pore residues of empty-pocket TRPV1 (orange) compared to apo TRPV1 (PDB-5IRZ). Density for M682 is shown. **(d)** Comparing the density for an outer leaflet lipid in the apo (PDB-5IRZ) and the empty-pocket states. The resident lipid in PDB-5IRZ (black sticks) and the outer leaflet lipid (dark red sticks) are shown superimposed beside the 25° C data to highlight the binding overlap of the two lipids. **(e)** Upper methionine restriction (M644) and G683 of apo TRPV1 (PDB-5IRZ). **(f)** Upper methionine restriction (M644) and lower methionine restriction (M682) of TRPV1 in the empty-pocket pocket at 25° C.

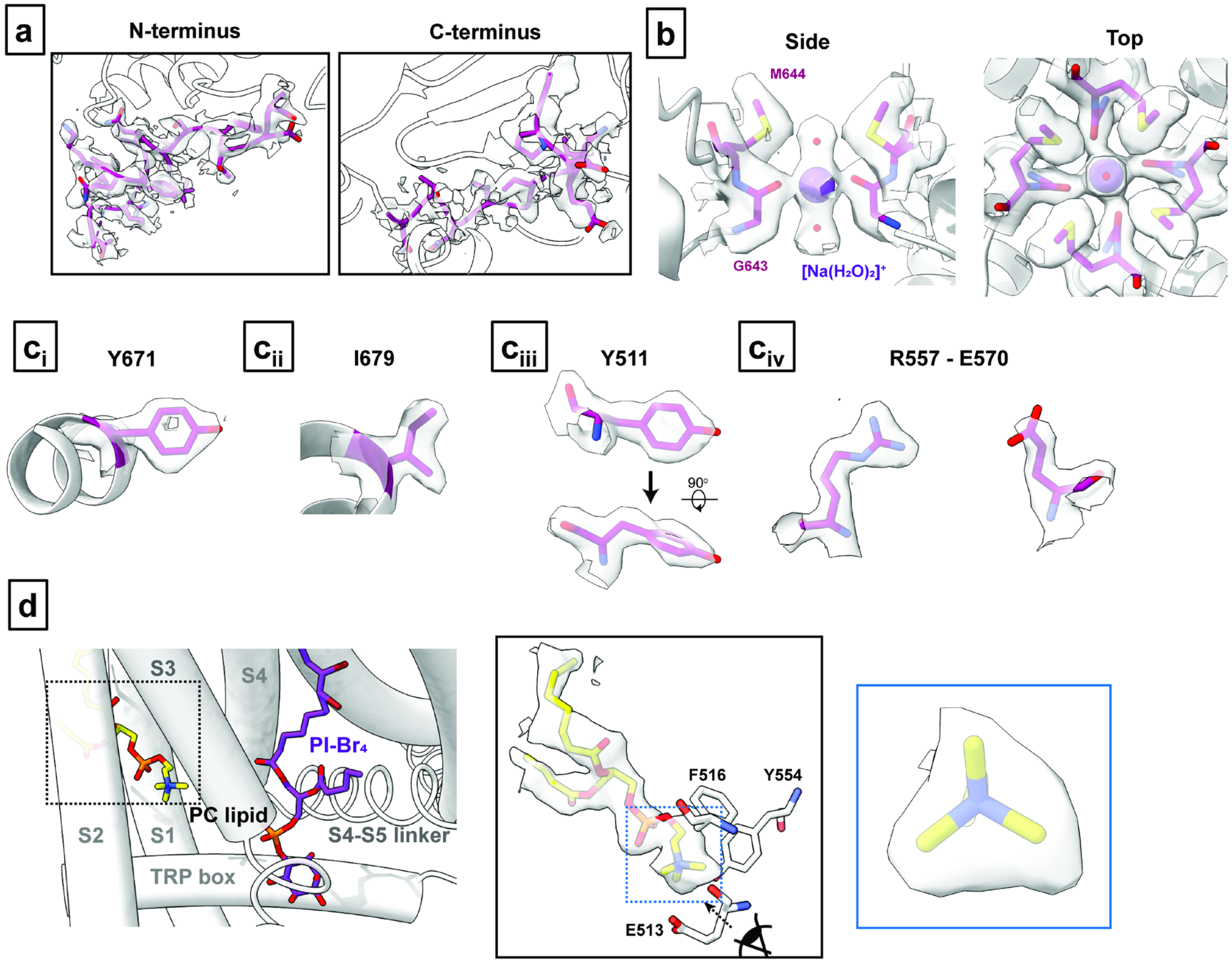


Supplementary Figure 2. *Structural details revealed by the PI-Br$_4$ data.* **(a)** Higher resolution for the N-terminus and C-terminus of the PI-Br$_4$ consensus map allows for further modeling of the N-terminus (starting at residue 177) and the remaining C-terminus of the truncated TRPV1 construct (up to residue 764). **(b)** Captured density at the selectivity filter in the PI-Br$_4$ consensus map models well to a $[Na(H_2O)_2]^+$ cation coordinated to G643. **(c)** Well-defined densities for key residues as revealed in the PIBr$_4$ consensus map. **(d)** Lipid density in a binding pocket within the S1 to S3 transmembrane helices shows well-defined features for a PC headgroup, importantly the tri-lobular of the quaternary amine group. Data shown is from the PI-Br$_4$ data in Conformation 1.

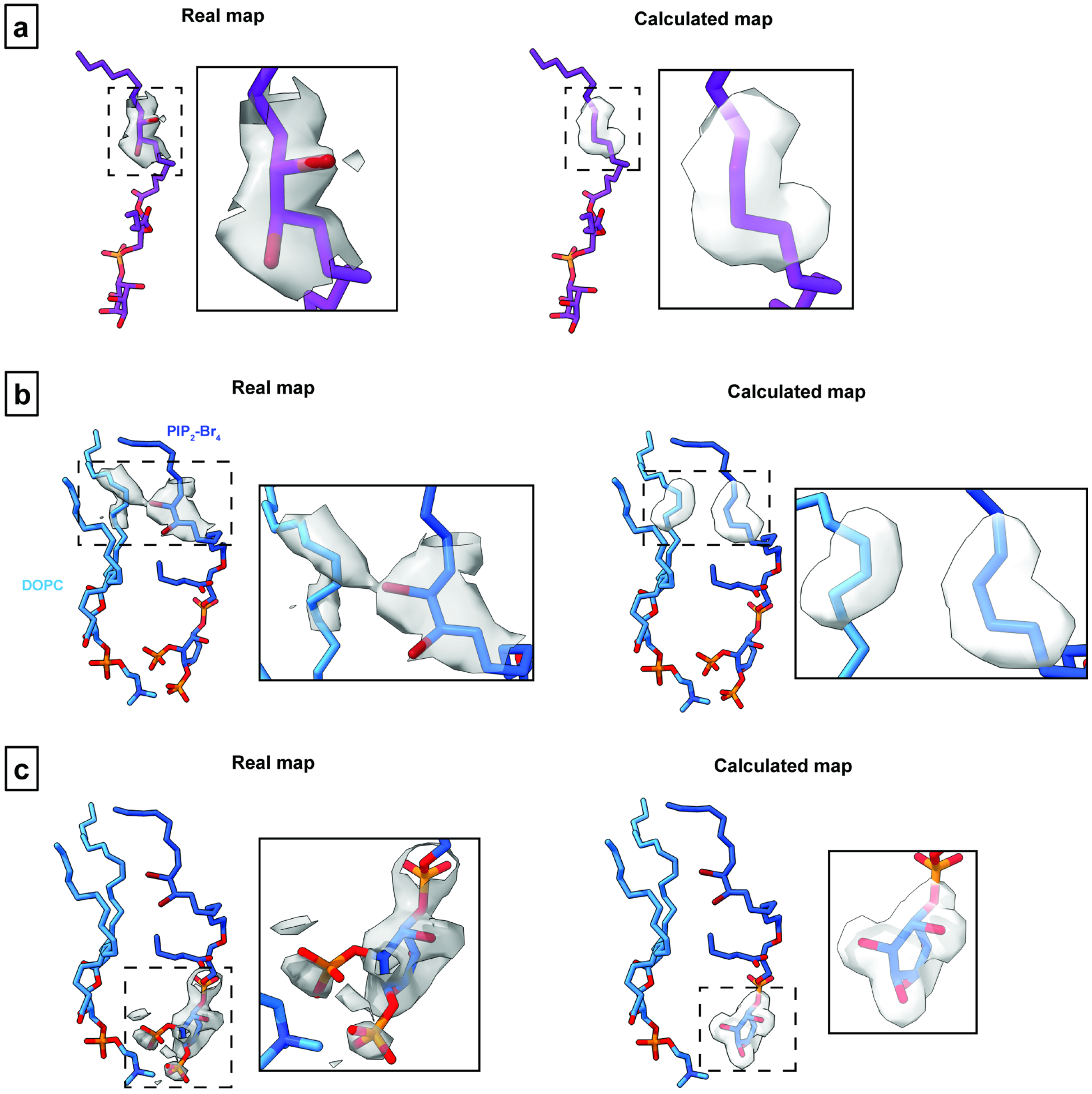


Supplementary Figure 3. *Real and calculated maps used for difference map analysis*. **(a)** Maps used for PI-$Br_4$ analysis. **(b)** Maps used for the bromine analysis for $PIP_2$-$Br_4$. **(c)** Maps used for the headgroup analysis for $PIP_2$-$Br_4$. Insets show close ups of the maps.

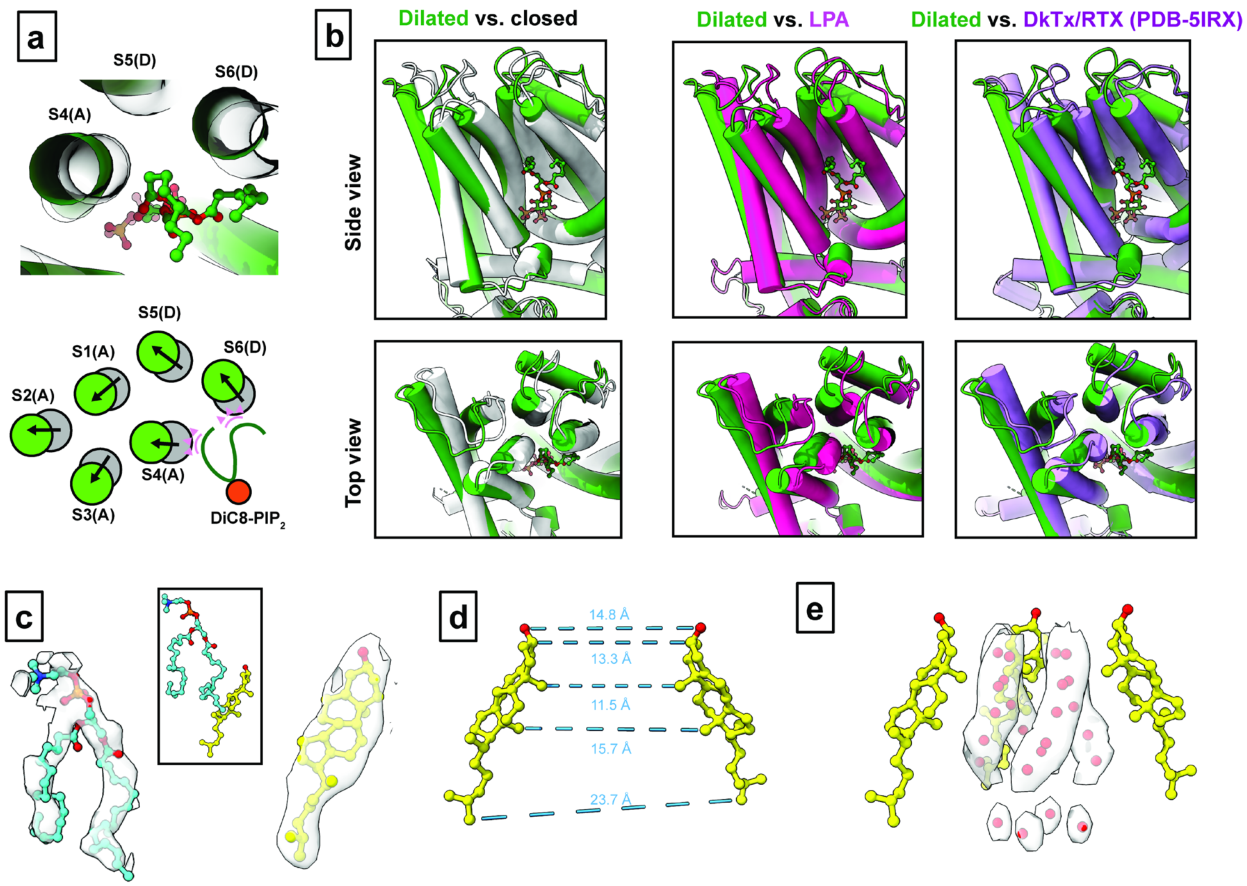


Supplementary Figure 4. *TRPV1 bound with diC8-PIP$_2$ in the dilated state.* **(a)** Top: Top-down view of diC8-PIP$_2$ (green sticks and balls) and surrounding helices. Grey ghost tubes are TRPV1 bound with diC8-PIP$_2$ in the closed state and green tubes are in the dilated state. Bottom: schematic comparing the helical positions between the closed and dilated states. Pink triangles indicate the steric overlap between the diC8-PIP$_2$ molecule in the dilated state and the transmembrane helices of TRPV1 in the closed state. **(b)** Comparisons of the dilated state of TRPV1 with diC8-PIP$_2$ bound and other states. **(c)** Densities for cholesterol and a lipid (modeled as DOPC) that is found in a cleft between the S6 and S5 helices of adjacent monomers (see Figure 3). **(d)** Interatom distances between two opposite cholesterol molecules demonstrating the pore radius formed by cholesterol. **(e)** Extra densities within the pore that are modeled as ordered water molecules due to the hydrophobic effect. 3 of the 4 cholesterol molecules are shown for clarity.

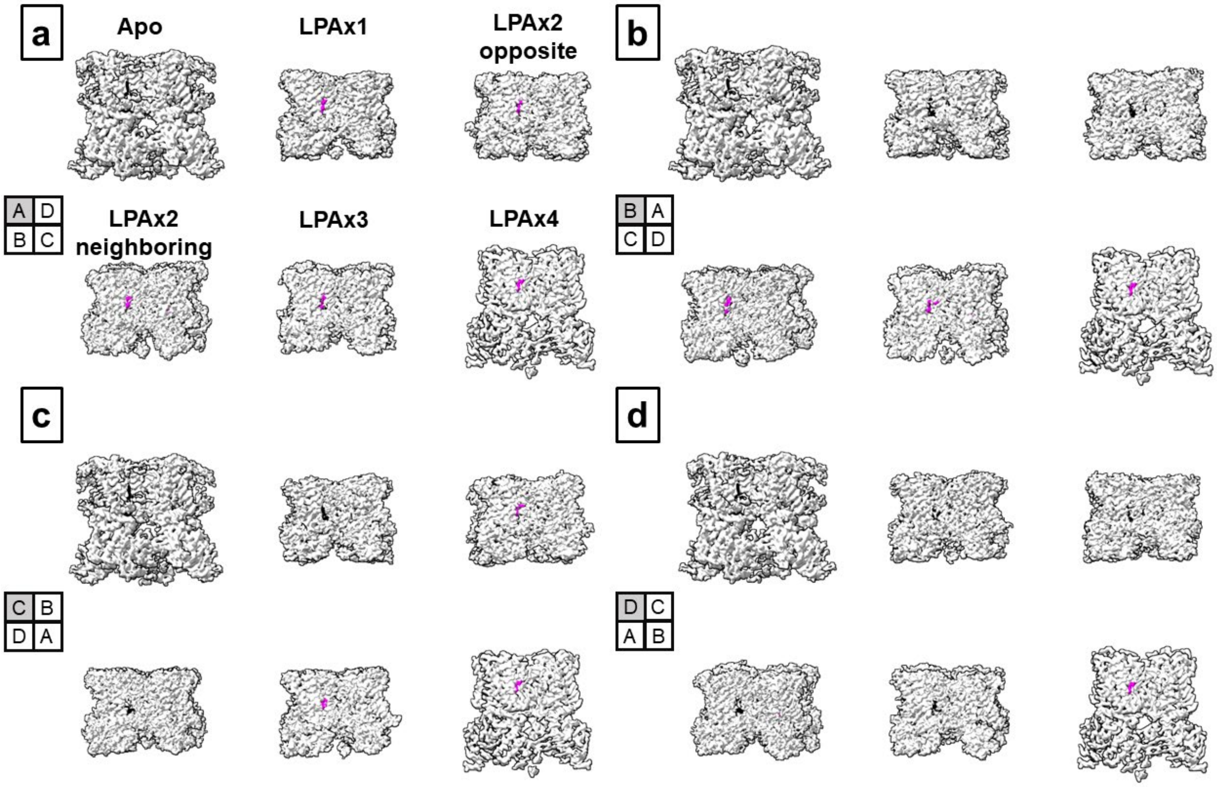


Supplementary Figure 5. *Density maps of TRPV1 bound with 0-4 LPA*. Each subpanel **(a-d)** shows TRPV1 rotated 90$^{o}$ counterclockwise starting with pocket A **(a)** and ending with pocket D **(d)**. Density corresponding to the resident phosphoinositide lipid is shown as black and density corresponding to LPA is shown as magenta.

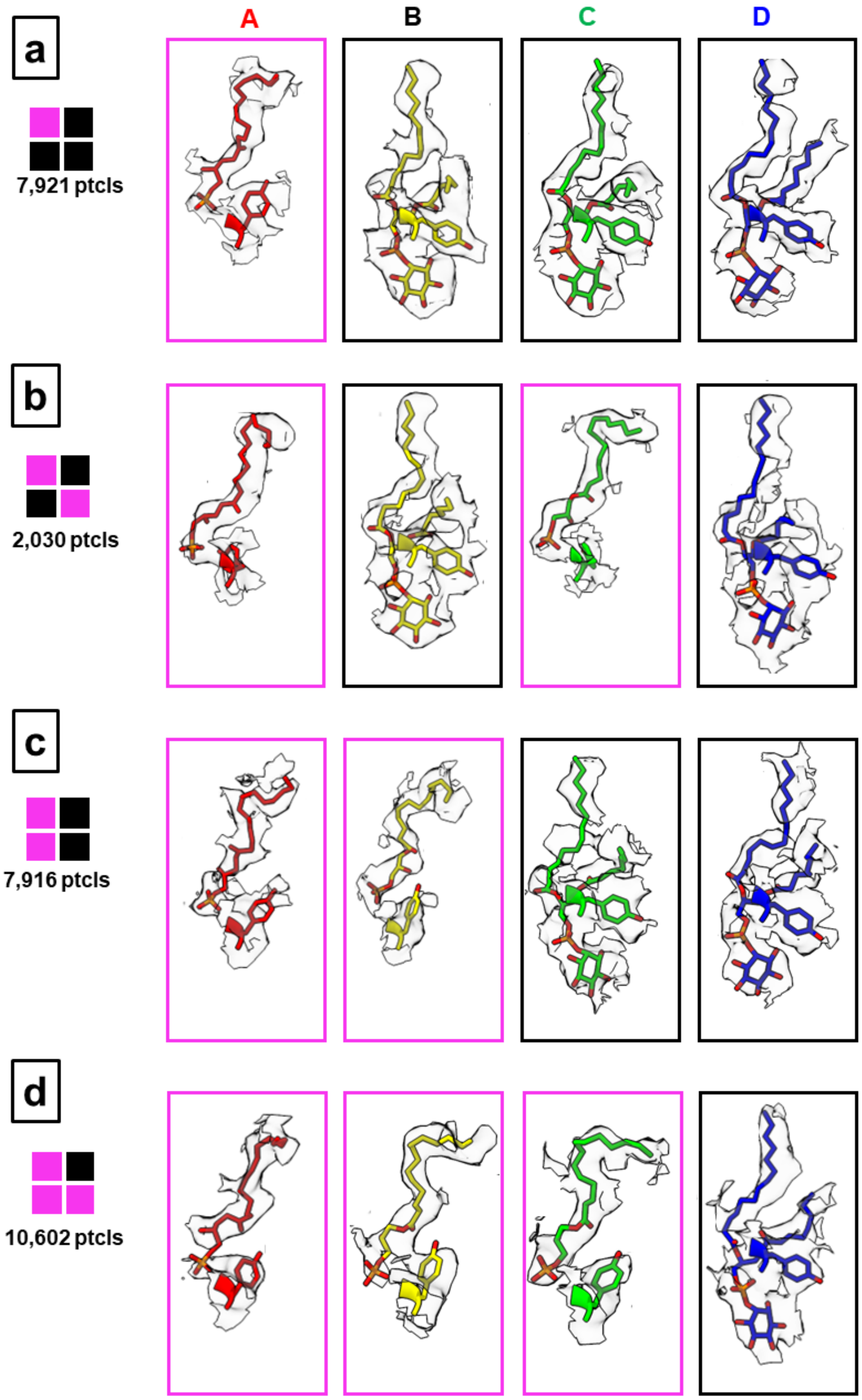


Supplementary Figure 6. *LPA sub-stoichiometric ligand densities.* Densities for Y511 and LPA or the resident lipid are shown for each monomer. LPA = magenta; resident lipid = black. **(a)** LPAx1. **(b)** LPAx2 opposite. **(c)** LPAx2 neighboring. **(d)** LPAx3.

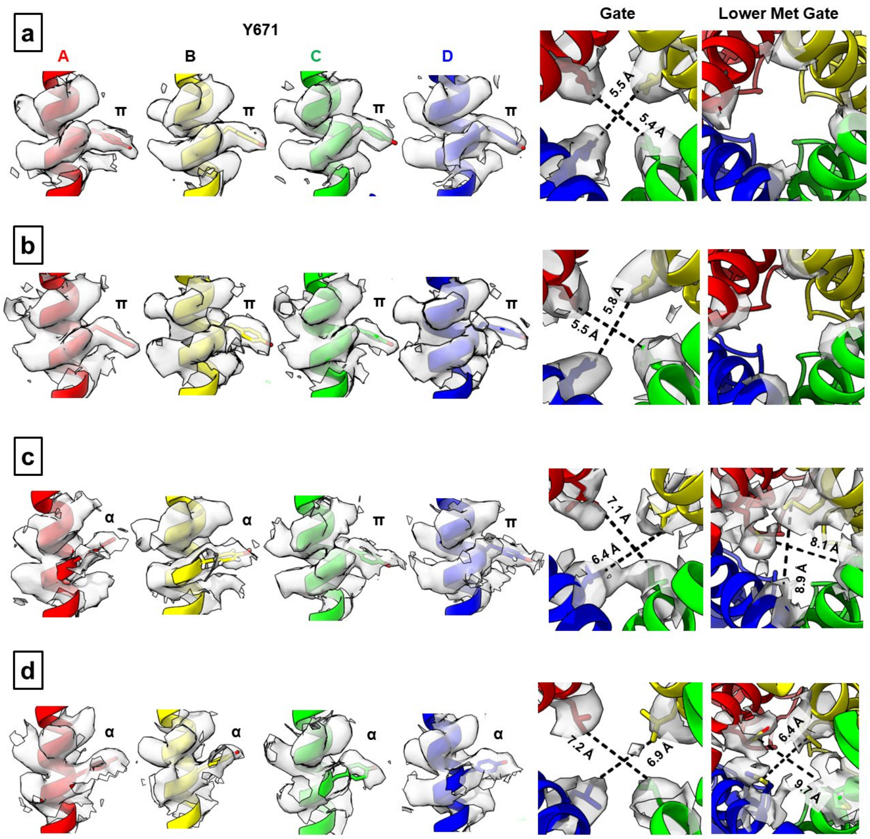


Supplementary Figure 7. *Densities of key residues lining the pore of the sub-saturating states.* **(a)** LPAx1. **(b)** LPAx2 in opposite pockets. **(c)** LPAx2 in neighboring pockets. **(d)** LPAx3. Interatom distances of the closest non-hydrogen atoms for the gating residues and the lower methionine filter are shown.

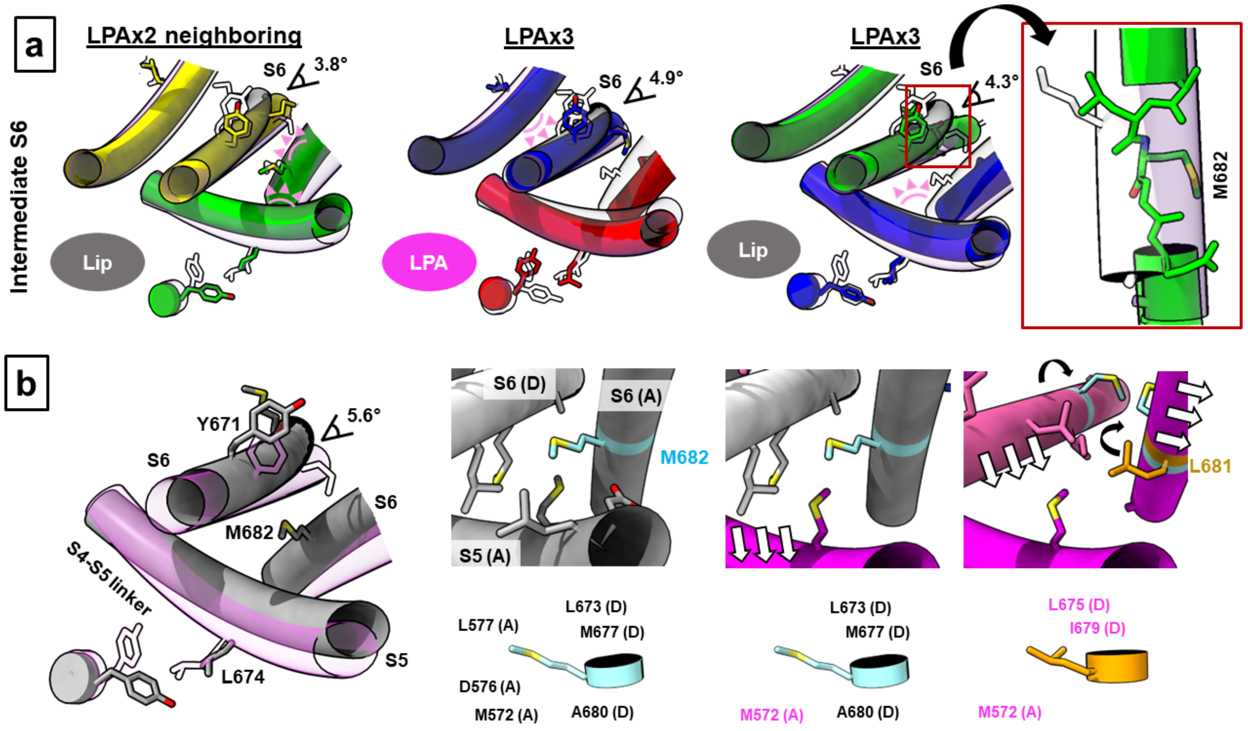


Supplementary Figure 8. *Intermediate states and mechanism of allostery* **(a)** Intermediate states of the S6 helices. Steric clashes preventing full opening of the S6 helix are shown as pink triangles. Angles of opening are shown as a measure of the intermediate state. For LPAx3, the S6-C helix is distorted from a typical α-helical structure. **(b)** Proposed mechanism for the LPA-induced opening of TRPV1 as the channel goes from apo state (grey) to one completely occupied by LPA (magenta). Residues in contact with M682 (< 5.0 Å) as it switches positions with L681 due to the π-α helix transition are highlighted. The S6 helix tilts by 5.6° during the transition from closed to open states.

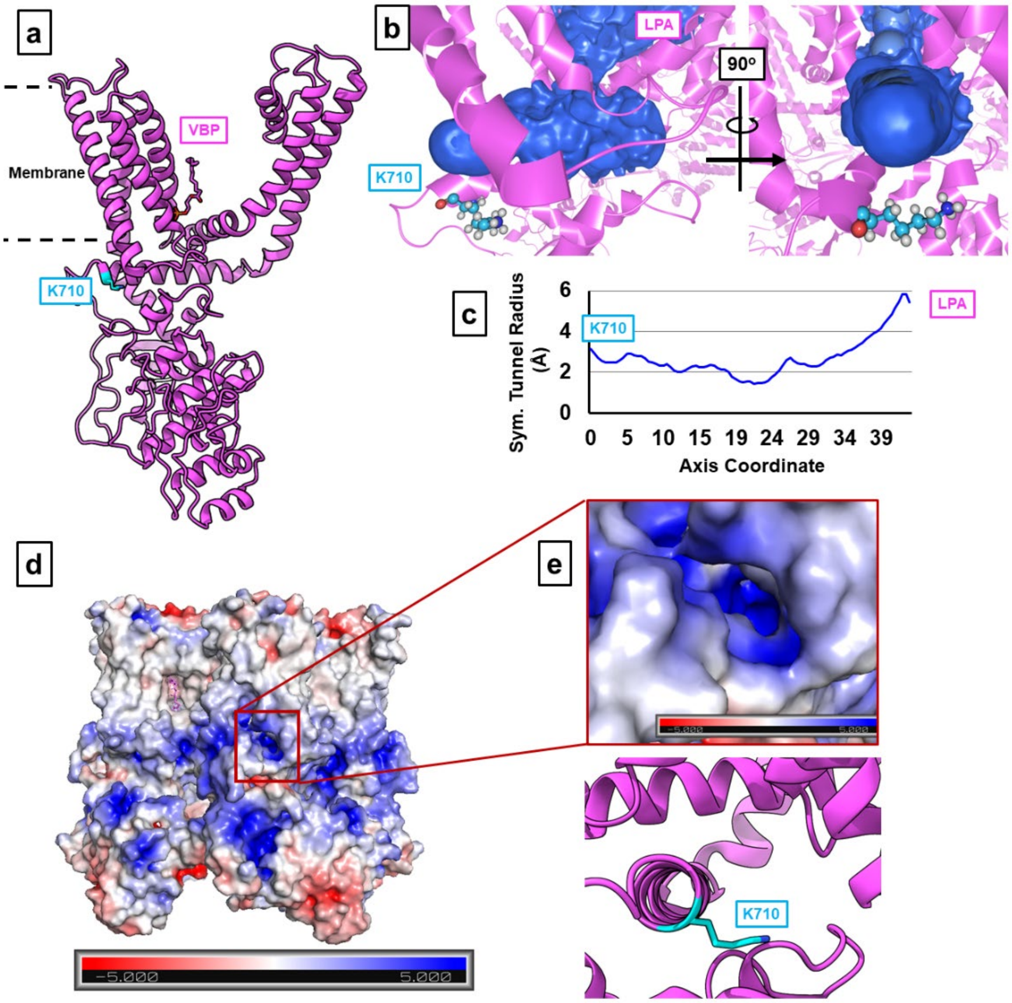


Supplementary Figure 9. *K710 marks a potential ingress tunnel for LPA.* **(a)** K710 (blue) sits near the membrane surface and approximately 20 Å away from the VBP. **(b)** An access tunnel (blue) connecting K710 to the VBP as identified by the program CAVER. **(c)** Size of the tunnel as determined from CAVER. **(d)** Surface charge map of TRPV1 with location of putative tunnel indicated by red box. (e) Surface charge map of the tunnel identified by CAVER with the corresponding model of TRPV1-LPAx4 below.

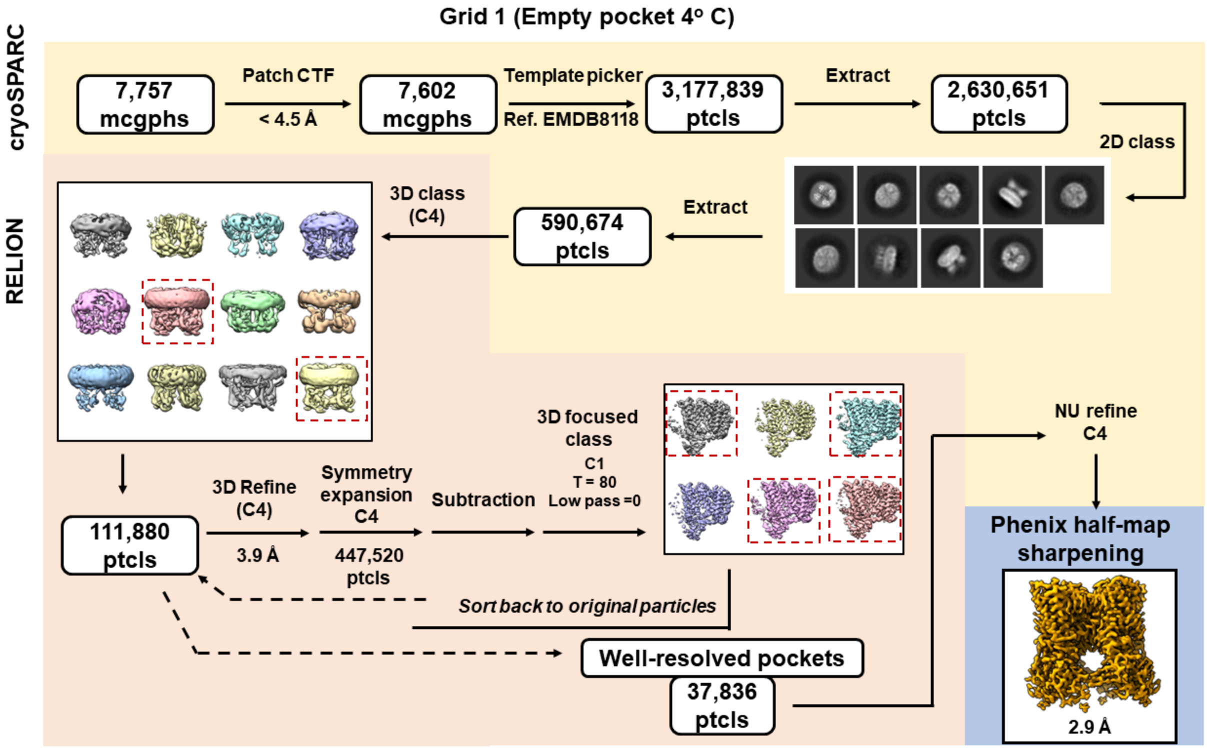


Supplementary Figure 10. Data processing scheme for Grid 1.

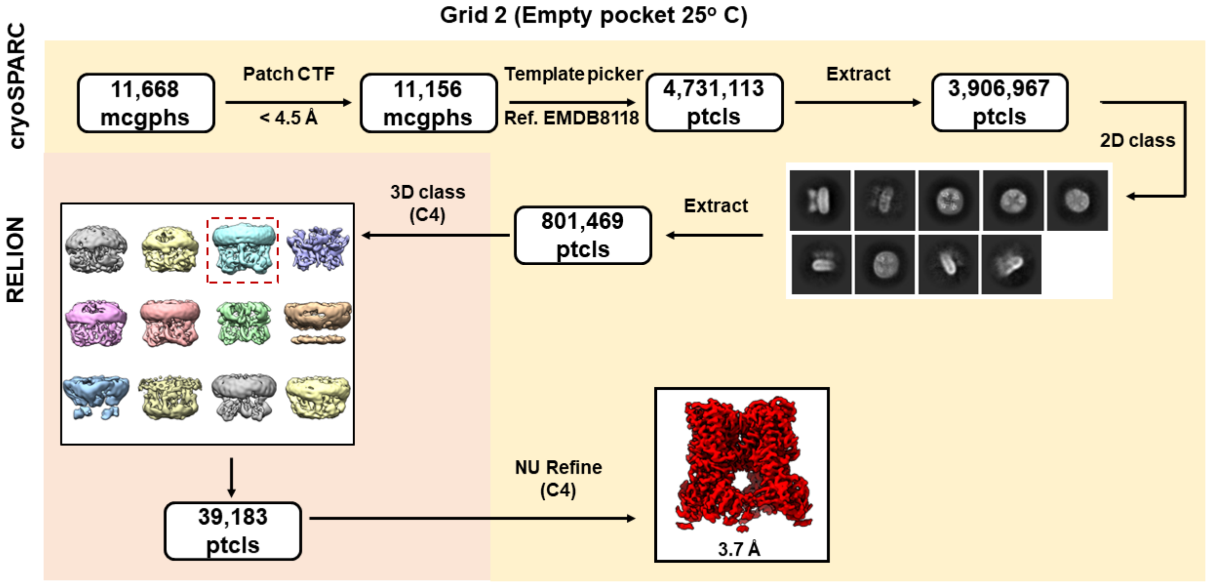


Supplementary Figure 11. Data processing scheme for Grid 2.

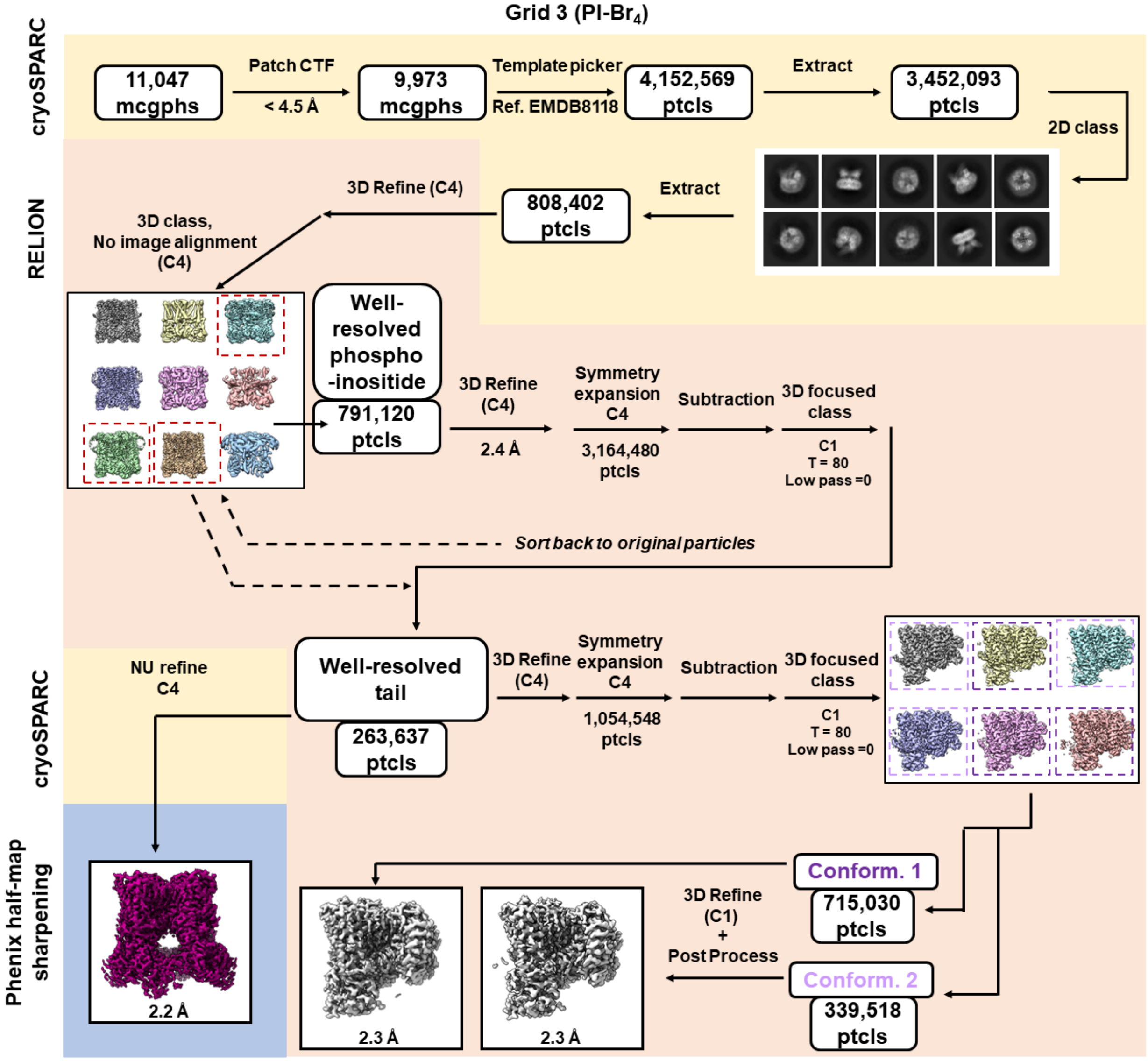


Supplementary Figure 12. Data processing scheme for Grid 3.

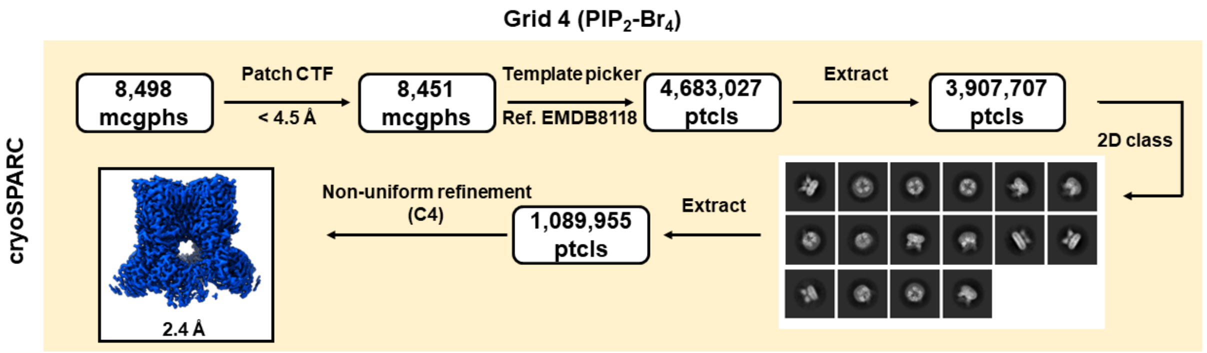


Supplementary Figure 13. Data processing scheme for Grid 4.

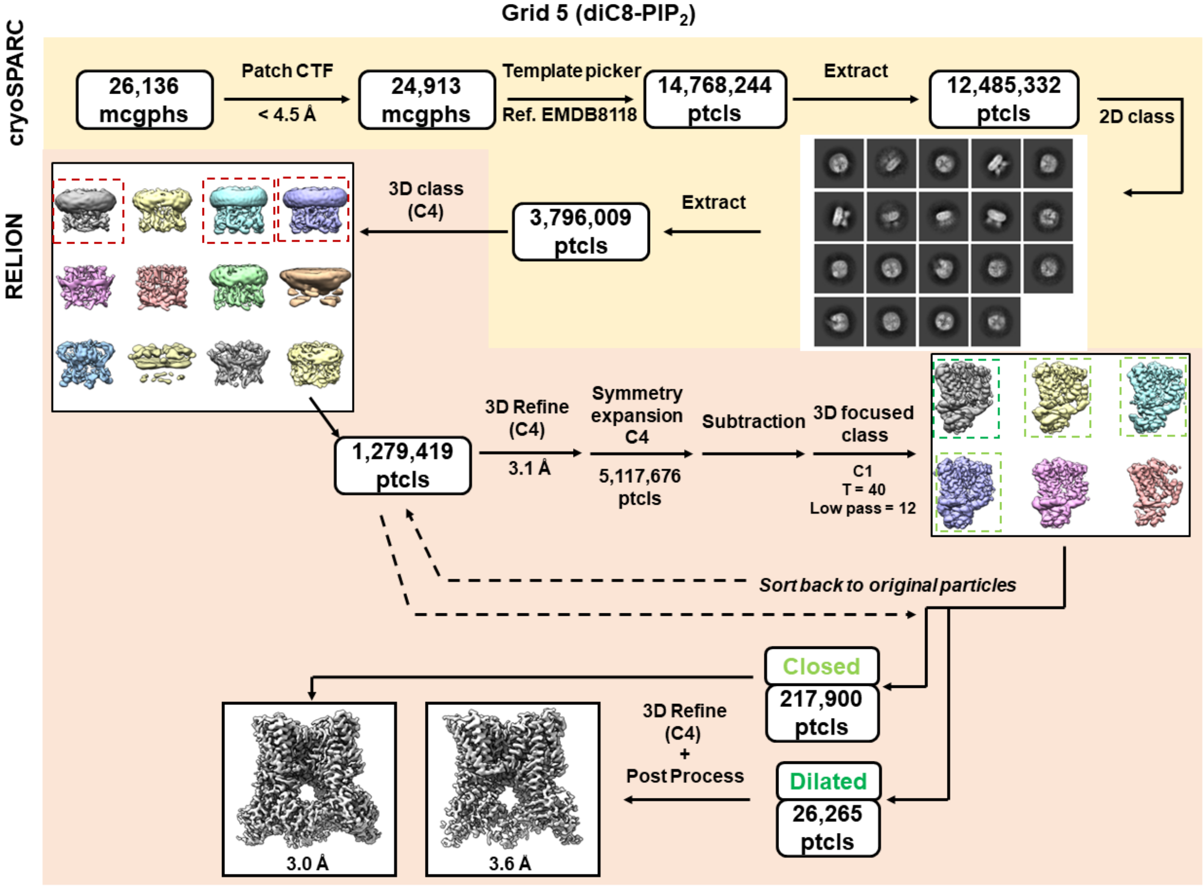


Supplementary Figure 14. Data processing scheme for Grid 5.

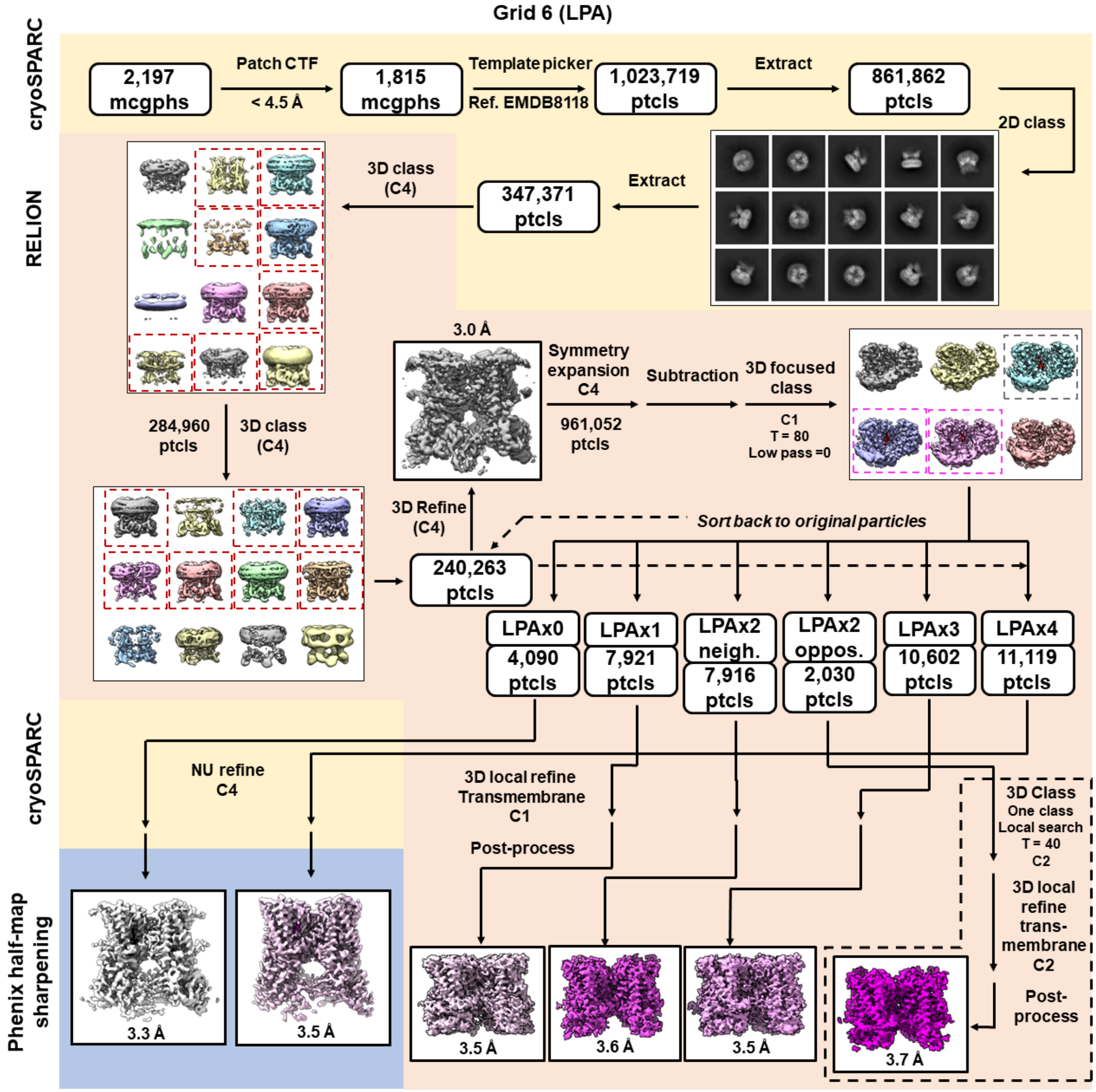


Supplementary Figure 15. Data processing scheme for Grid 6.

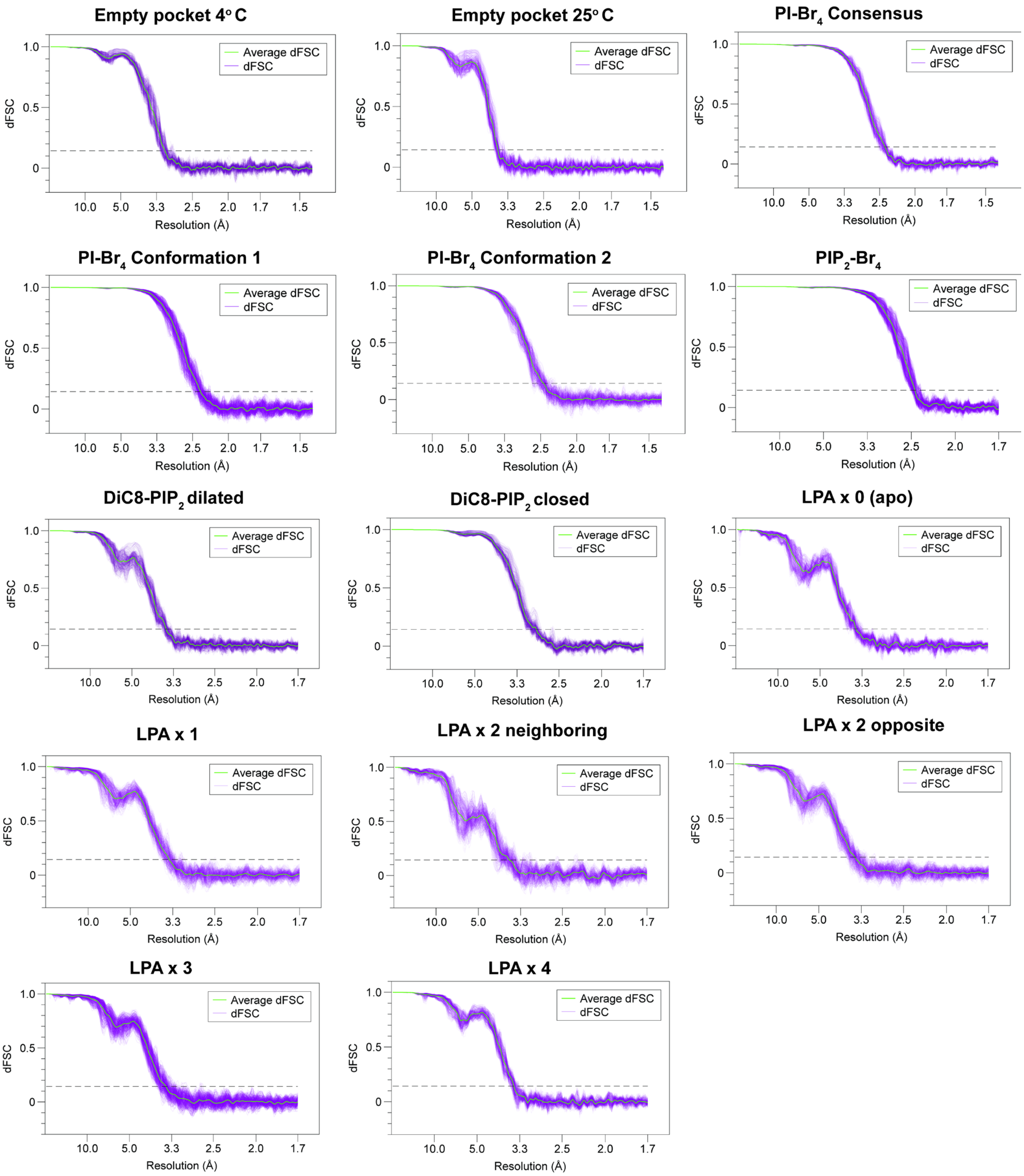


Supplementary Figure 16. Plots of the dFSC curves for maps in the paper. Gold standard FSC (0.143) is shown as a dashed line.

Supplementary Table 1. Data acquisition (Grids 1 and 2).

| | | Empty Pocket 4ºC (Grid 1) | Empty Pocket 25ºC (Grid 2) |
|---|---|---|---|
| **Ascension Codes** | | | |
| | PDB | 8U3J | 8U3L |
| | EMD | 41864 | 41866 |
| **Atoms** | | | |
| | All | 35,217 | 35,217 |
| | Hydrogens | 17,724 | 17,724 |
| | Residues | 2,136 | 2,136 |
| **Ligands** | | | |
| | Na+ | 1 | 1 |
| | Phosphatidic acid (3PH) | 1 | 1 |
| **RMSZ deviation** | | | |
| | bond length (Å) | 0.28 | 0.28 |
| | bond angle (°) | 0.50 | 0.47 |
| **Validation** | | | |
| | Clash score | 5 | 5 |
| | Poor rotamers (%) | 0 | 0 |
| **Ramachandran** | | | |
| | Favored (%) | 95 | 94 |
| | Allowed | 5 | 6 |
| | disallowed | 0 | 0 |
| **Microscope** | | | |
| | Microscope ID | TEM gamma ($S^2C^2$) | TEM beta ($S^2C^2$) |
| | Magnification | 130,000 × | 130,000 × |
| | Voltage (KV) | 300 | 300 |
| | Dose ($e^-$/pix/frame) | 1.02 | 1.02 |
| | Total dose ($e^-$/Å) | 60 | 60 |
| | Defocus range (μm) | 0.5-2.0 | 0.5-2.0 |
| | Pixel size (Å) | 0.68 | 0.68 |
| **Data processing** | | | |
| | micrographs | 7,757 | 11,668 |
| | initial particles | 590,647 | 801,469 |
| | final particles | 37,836 | 39,183 |
| | Box size (pix) | 416 | 416 |
| | Symmetry | C4 | C4 |
| | Map resolution (Å) | 2.9 | 3.7 |
| | FSC threshold | 0.143 | 0.143 |
| | Map sharpening | Phenix Half-Map | CryoSPARC |
| | b-factor | NA | -152.4 |
| | Final adjusted map surface area | 1.98 | NA |
| | final map kurtosis | 128.61 | NA |

Supplementary Table 2. Data acquisition (Grids 3 and 4).

| | | PI-$Br_4$ consensus (Grid 3) | PI-$Br_4$ Conformation 1 (Grid 3) | PI-$Br_4$ Conformation 2 (Grid 3) | $PIP_2$-$Br_4$ (Grid 4) |
|---|---|---|---|---|---|
| **Ascension Codes** | | | | | |
| | PDB | 8U4D | 8U3A | 8U3C | 8U43 |
| | EMD | 41879 | 41855 | 41857 | 41873 |
| **Atoms** | | | | | |
| | All | 36,769 | 6,973 | 7,037 | 37,219 |
| | Hydrogens | 18,458 | 3,542 | 3,578 | 18,628 |
| | Residues | 2,236 | 405 | 405 | 2,224 |
| **Ligands** | | | | | |
| | Na+ | 1 | 1 | 1 | 1 |
| | PI-$Br_4$ (VPN) | 4 | 1 | 1 | 0 |
| | $PIP_2$-$Br_4$ (V5H) | 0 | 0 | 0 | 4 |
| | Phosphatidic acid (3PH) | 0 | 0 | 1 | 0 |
| | DOPC (PCW) | 0 | 1 | 1 | 4 |
| **RMSZ deviation** | | | | | |
| | bond length (Å) | 0.29 | 0.38 | 0.39 | 0.32 |
| | bond angle (°) | 0.49 | 0.55 | 0.56 | 0.55 |
| **Validation** | | | | | |
| | Clash score | 2 | 1 | 1 | 4 |
| | Poor rotamers (%) | 5 | 0 | 0 | 4 |
| **Ramachandran** | | | | | |
| | Favored (%) | 94 | 94 | 95 | 94 |
| | Allowed | 6 | 6 | 5 | 6 |
| | disallowed | 0 | 0 | 0 | 0 |
| **Microscope** | | | | | |
| | Microscope ID | UCSF | | | UCSF |
| | Magnification | 130,000 × | | | 105,000 × |
| | Voltage (KV) | 300 | | | 300 |
| | Dose rate ($e^-$/pix/s) | 16 | | | 16 |
| | Dose ($e^-$/pix/frame) | 0.54 | | | 0.57 |
| | Total dose ($e^-$/Å) | 47.2 | | | 45.8 |
| | Defocus range (µm) | 0.5-2.0 | | | 0.5-2.0 |
| | Pixel size (Å) | 0.644 | | | 0.835 |
| **Data processing** | | | | | |
| | micrographs | 11,047 | | | 8,498 |
| | initial particles | 808,402 | | | 1,089,955 |
| | final particles | 263,637 | 715,030 | 339,518 | 1,089,955 |
| | Box size (pix) | 416 | | | 416 |
| | Symmetry | C4 | C1 | C1 | C4 |
| | Map resolution (Å) | 2.2 | 2.3 | 2.3 | 2.4 |
| | FSC threshold | 0.143 | | | 0.143 |
| | Map sharpening | Phenix Half-Map | RELION | RELION | CryoSPARC |
| | b-factor | NA | -70.5 | -69.9 | -114.7 |
| | Final adjusted map surface area | -3.28 | NA | NA | NA |
| | final map kurtosis | 181.58 | NA | NA | NA |

Supplementary Table 3. Data acquisition (Grid5).

| | | DiC8-$PIP_2$ closed | DiC8-$PIP_2$ dilated |
|---|---|---|---|
| **Ascension Codes** | | | |
| | PDB | 8U30 | 8U2Z |
| | EMD | 41848 | 41847 |
| **Atoms** | | | |
| | All | 34,793 | 34,361 |
| | Hydrogens | 17,452 | 17,316 |
| | Residues | 2,112 | 2,040 |
| **Ligands** | | | |
| | Na+ | 1 | 1 |
| | DiC8-$PIP_2$ (PIO) | 4 | 4 |
| | Cholesterol (CLR) | 0 | 4 |
| | DOPC (PCW) | 0 | 4 |
| **RMSZ deviation** | | | |
| | bond length (Å) | 0.28 | 0.30 |
| | bond angle (°) | 0.48 | 0.55 |
| **Validation** | | | |
| | Clash score | 6 | 8 |
| | Poor rotamers (%) | 0 | 1 |
| **Ramachandran** | | | |
| | Favored (%) | 94 | 92 |
| | Allowed | 6 | 8 |
| | disallowed | 0 | 0 |
| **Microscope** | | | |
| | Microscope ID | UCSF | |
| | Magnification | 105,000 × | |
| | Voltage (KV) | 300 | |
| | Dose rate ($e^-$/pix/s) | 16 | |
| | Dose ($e^-$/pix/frame) | 0.57 | |
| | Total dose ($e^-$/Å) | 45.8 | |
| | Defocus range (µm) | 0.5-2.0 | |
| | Pixel size (Å) | 0.835 | |
| **Data processing** | | | |
| | micrographs | 26,136 | |
| | initial particles | 3,796,009 | |
| | final particles | 217,900 | 26,265 |
| | Box size (pix) | 384 | |
| | Symmetry | C4 | |
| | Map resolution (Å) | | 3.6 |
| | FSC threshold | 0.143 | |
| | Map sharpening | RELION | |
| | b-factor | -112.4 | -113.5 |

Supplementary Table 4. Data acquisition (Grid 6).

| | | LPAx0 | LPAx1 | LPAx2 opposite | LPAx2 neighboring | LPAx3 | LPAx4 |
|---|---|---|---|---|---|---|---|
| **Ascension Codes** | | | | | | | |
| | PDB | 8T0E | 8T0Y | 8T10 | 8T3L | 8T3M | 8T0C |
| | EMD | 40941 | 40949 | 40951 | 41005 | 41006 | 40940 |
| **Atoms** | | | | | | | |
| | All | 35,109 | 21,244 | 20,332 | 20,300 | 22,473 | 34,994 |
| | Hydrogens | 17,676 | 10,635 | 10,250 | 10,234 | 11,317 | 17,596 |
| | Residues | 2,124 | 1,273 | 1,198 | 1,197 | 1,345 | 2,128 |
| **Ligands** | | | | | | | |
| | Na+ | 1 | 1 | 1 | 2 | 2 | 2 |
| | Resident lipid (85R) | 4 | 3 | 2 | 2 | 1 | 4 |
| | LPA (NKN) | 0 | 1 | 2 | 2 | 3 | 0 |
| **RMSZ deviation** | | | | | | | |
| | bond length (Å) | 0.37 | 0.32 | 0.46 | 0.32 | 0.34 | 0.28 |
| | bond angle (°) | 0.55 | 0.48 | 0.60 | 0.49 | 0.50 | 0.48 |
| **Validation** | | | | | | | |
| | Clash score | 5 | 4 | 4 | 5 | 7 | 5 |
| | Poor rotamers (%) | 2 | 1 | 3 | 1 | 1 | 0 |
| **Ramachandran** | | | | | | | |
| | Favored (%) | 90 | 93 | 94 | 93 | 92 | 94 |
| | Allowed | 10 | 7.0 | 6.0 | 7.0 | 8.0 | 6.0 |
| | disallowed | 0 | 0 | 0 | 0 | 0 | 0 |
| **Microscope** | | | | | | | |
| | Microscope ID | UCSF | | | | | |
| | Magnification | 105,000 × | | | | | |
| | Voltage (KV) | 300 | | | | | |
| | Dose rate ($e^-$/pix/s) | 16 | | | | | |
| | Dose ($e^-$/pix/frame) | 0.57 | | | | | |
| | Total dose ($e^-$/Å) | 45.8 | | | | | |
| | Defocus range (μm) | 0.5-2.0 | | | | | |
| | Pixel size (Å) | 0.835 | | | | | |
| **Data processing** | | | | | | | |
| | micrographs | 1,815 | | | | | |
| | initial particles | 240,263 | | | | | |
| | final particles | 4,090 | 7,921 | 2,030 | 7,916 | 10,602 | 11,119 |
| | Box size (pix) | 384 | | | | | |
| | Symmetry | C4 | C1 | C2 | C1 | C1 | C4 |
| | Map resolution (Å) | 3.3 | 3.5 | 3.7 | 3.6 | 3.5 | 3.5 |
| | FSC threshold | 0.143 | | | | | |
| | Map sharpening | Phenix half-map | RELION | RELION | RELION | RELION | Phenix Half-map |
| | b-factor | NA | -76.3 | -79.9 | -75.1 | -90.1 | NA |
| | Final adjusted map surface area | 2.05 | NA | NA | NA | NA | 1.91 |
| | final map kurtosis | 94.36 | NA | NA | NA | NA | 122.67 |
